# Human immune phenotyping reveals accelerated aging in type 1 diabetes

**DOI:** 10.1101/2023.02.24.529902

**Authors:** Melanie R. Shapiro, Xiaoru Dong, Daniel J. Perry, James M. McNichols, Puchong Thirawatananond, Amanda L. Posgai, Leeana Peters, Keshav Motwani, Richard S. Musca, Andrew Muir, Patrick Concannon, Laura M. Jacobsen, Clayton E. Mathews, Clive H. Wasserfall, Michael J. Haller, Desmond A. Schatz, Mark A. Atkinson, Maigan A. Brusko, Rhonda L. Bacher, Todd M. Brusko

## Abstract

The composition of immune cells in peripheral blood is dramatically remodeled throughout the human lifespan, as environmental exposures shape the proportion and phenotype of cellular subsets. These dynamic shifts complicate efforts to identify disease-associated immune signatures in type 1 diabetes (T1D), which is variable in age of onset and rate of β-cell decline. Herein, we conducted standardized flow cytometric immune profiling on peripheral blood from a cross-sectional cohort of T1D participants (*n*=240), their first-degree relatives (REL, *n*=310), those at increased risk with two or more islet autoantibodies (RSK, *n*=24), and autoantibody negative healthy controls (CTR, *n*=252). We constructed an immune-age predictive model in healthy subjects and developed an interactive data visualization portal (ImmScape; https://ufdiabetes.shinyapps.io/ImmScape/). When applied to the T1D cohort, this model revealed accelerated immune aging (p<0.001) as well as phenotypic signatures of disease after age correction. Of 192 investigated flow cytometry and complete blood count readouts, 46 were significantly associated with age only, 25 with T1D only, and 23 with both age and T1D. Phenotypes associated with T1D after age-correction were predictive of T1D status (AUROC=82.3%). Phenotypes associated with accelerated aging in T1D included increased CXCR3^+^ and PD-1^+^ frequencies in naïve and memory T cell subsets, despite reduced PD-1 expression levels (mean fluorescence intensity) on memory T cells. Additionally, quantitative trait locus analysis linked an increase in HLA-DR expression on monocytes with the T1D-associated HLA-DR4/DQ8 genotype, regardless of clinical group. Our findings demonstrate advanced immune aging in T1D and highlight disease-associated phenotypes for biomarker monitoring and therapeutic interventions.

**One Sentence Summary:** Peripheral blood characterization reveals accelerated immune-age and age-adjusted proinflammatory immune phenotypes in type 1 diabetes.

## INTRODUCTION

With improved diabetes classification tools, it is now appreciated that the onset of type 1 diabetes (T1D) may occur throughout the human lifespan (1; 2), although the majority of cases develop within two peaks between ages five to seven years or near puberty (3). Individuals with high genetic risk for T1D are more likely to receive diagnoses during these peak time periods (4; 5), and have been shown to demonstrate the appearance of islet cell-reactive autoantibodies (AAb) indicative of disease progression as early as within the first two years of age (6; 7). The presence of an islet AAb precedes insulitis, the appearance of immune cell infiltrates in pancreatic islets. The composition of insulitis has been demonstrated to depend upon the age-at-onset of T1D, which is largely driven by HLA risk (5; 8). CD20^+^ B cells constitute a major proportion of the pancreatic immune infiltrate in individuals below the age of seven years, whereas those above age thirteen possess a more CD8^+^, CD4^+^, and CD68^+^ cell-dominated insulitis (9; 10), suggesting that the immune populations involved in disease pathogenesis vary by age at diagnosis.

The quest for cellular biomarkers of disease pathogenesis in T1D is limited to recirculating immune cells that do not perfectly reflect the activity at the priming lymph nodes and autoimmune lesion (11), and these challenges are further compounded by variations in peripheral blood immune cell subset composition due to age and environmental exposures as major drivers over heritable factors (12). To address the problem of confounding technical and biological factors masking disease-related changes in the immune system, the Human Immunophenotyping Consortium (HIPC) has developed a recommended set of flow cytometry panels to allow for aggregation and comparison of data across studies, facilitating improved matching of data from healthy controls to those with disease (13). The HIPC panels were designed to quantify memory T cell, regulatory T cell (Treg), effector T cell (Teff), B cell, dendritic cell (DC), monocyte, and natural killer (NK) cell subset proportions and phenotypes (13). These standardized panels have been successfully used to identify immune modulation due to vaccination (14; 15), infection (16; 17), autoimmunity (18; 19), and cancer (20; 21), although full HIPC phenotyping of T1D has not been previously performed.

The impact of aging on immune phenotypes is well-established (22; 23), wherein environmental exposures, particularly infection by cytomegalovirus (CMV), are known to drive expansion of a large pool of antigen-specific T cells that persist in circulation with a memory phenotype (24; 25). However, these previous investigations have primarily been limited to cohorts of adults. Here, we studied a more extensive age range (2-83 years) in order to capture the immune dynamics of childhood and adolescence, during which key events such as vaccination and infection may alter immune maturation at differing rates compared to adults. In order to study disease-mediated perturbations of the immune system, our cross-sectional cohort (*n*=826) was designed to include healthy controls (CTR, *n*=252) and individuals across the range of risk for T1D progression: unaffected first-degree relatives of individuals with T1D (REL, *n*=310), rare at-risk participants who have two or more islet AAb (RSK, *n*=24), and those diagnosed with T1D (*n*=240).

In this study, we categorized major patterns of immune subset trajectories in healthy individuals over the pediatric and adult age range, allowing for detection of modulated trajectories in T1D. We further modeled the immunophenotyping data to estimate immunological age (26) as compared to chronological age in individuals with T1D versus those without diabetes. Age-corrected individual phenotypes contributing to differences in immunological age were compared across all participants binned by progressive T1D risk or status (27) to understand whether each was likely to contribute to immune activation and disease pathogenesis as opposed to a consequence post-onset (e.g., dysglycemia-induced inflammation (28)). Lastly, we assessed associations between T1D genetic risk (29–31) and immunophenotypes, the results of which supported the notion that the majority of observed immune perturbations in T1D reflect accelerated immune aging as a feature of disease pathogenesis. Notably, all immunophenotyping data generated herein are available for visualization and analysis via an interactive R/Shiny application (ImmScape; https://ufdiabetes.shinyapps.io/ImmScape/).

## RESULTS

### Impact of age on immune population dynamics

From rested peripheral blood samples, we used six flow cytometry panels adapted from HIPC recommendations (13) as we previously published (18; 32), to generate detailed immunophenotyping data encompassing proportions of innate and adaptive immune cells as well as phenotypes of memory T cell, Treg, Teff, T follicular helper (Tfh), B cell, DC, monocyte, and NK cell subsets (**Figure 1A**, with detailed gating schematics in **Figures S1-S6**). We first tested an initial cohort of 12 individuals in duplicate to evaluate assay reproducibility and observed that the biological coefficient of variance (CV) largely outweighed variance between technical duplicates (45.23 ± 25.66% vs. 8.87 ± 7.56%, **Figure 1B**), in agreement with previously established guidelines for replicability in flow cytometric studies (33). These studies were then extended to characterize flow cytometric immunophenotypes with accompanying complete blood count (CBC) measurements, for a total of 192 total outcome measures, on a large cross-sectional cohort of *n*=826 persons with or at varying levels of risk for T1D (**Table 1**). The majority of CBC values were within normal range with the following exceptions: low mean corpuscular hemoglobin concentration (MCHC), low neutrophil percentage, and high lymphocyte percentage were observed across CTR, REL, and T1D groups, presumably related to transport and storage time; T1D subjects displayed increased hematocrit percentage and increased platelet count, potentially reflecting dehydration and hyperglycemia, respectively (34) (**Table S1**). Covariates including age, sex, BMI percentile, and race differed between cohorts with varying levels of T1D risk (27) (**Table 1**). The age distribution between each group differed quite significantly, where the AAb- and T1D cohorts both have bimodal age distributions with different proportions in each component, as expected in a mostly pediatric T1D cohort with unaffected individuals being comprised largely of siblings and parents attending clinic visits. Upon noting this age discordance, we initially quantified how each immune phenotype changed with age in unaffected individuals consisting of islet AAb negative (AAb-) CTR and islet AAb-REL. We performed Spearman correlation analyses between age and peripheral blood phenotypes (obtained by CBC and flow cytometry), demonstrating that the majority of age-related associations appeared in the adaptive compartment as opposed to innate cells or CBC parameters (**Figure 1C-D**). In support of this notion, visual inspection of the dynamics of major subsets defined by our flow cytometry panels revealed relatively consistent proportions of innate cells across age, including monocytes, DCs, and NK cells, in contrast to adaptive immune subsets within the B cell and T cell compartments, which showed distinct age-related contraction of naïve and expansion of memory subsets (**Figure 1E-L**).

**Fig. 1.**
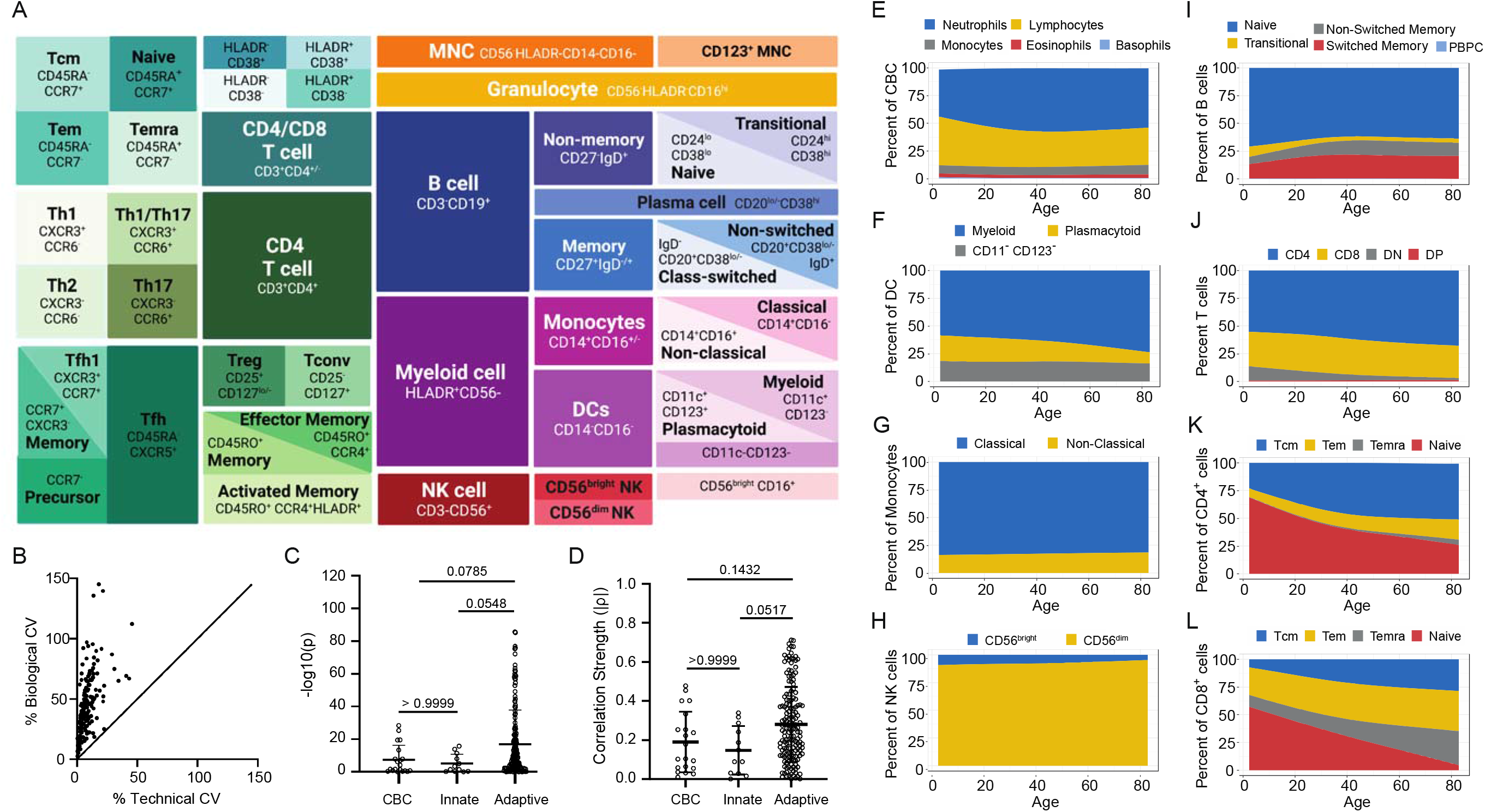
Immune population dynamics and QC of outcome measures. (**A**) Schematic representation of hierarchal gating strategy used to identify 172 immune cell subsets evaluated from human peripheral blood. An additional 20 parameters were derived from CBC. (**B**) Low technical variation observed from peripheral blood samples (*n*=12) stained in duplicate for assessment by flow cytometry. (**C**) -log10(p) (**D**) and correlation strength (absolute value of ρ) from Spearman correlation between phenotypes (each phenotype is a data point) and age shows strongest associations in the adaptive compartment as compared to innate or complete blood count (CBC). Kruskal-Wallis test with Dunn’s multiple comparisons test results denoted above bars. (**E-L**) Phenotype proportions estimated using a smoothing spline model as a function of age in AAb-individuals. (See also Figures S1-S6, Table S1 and Table S6).

**Table 1.**
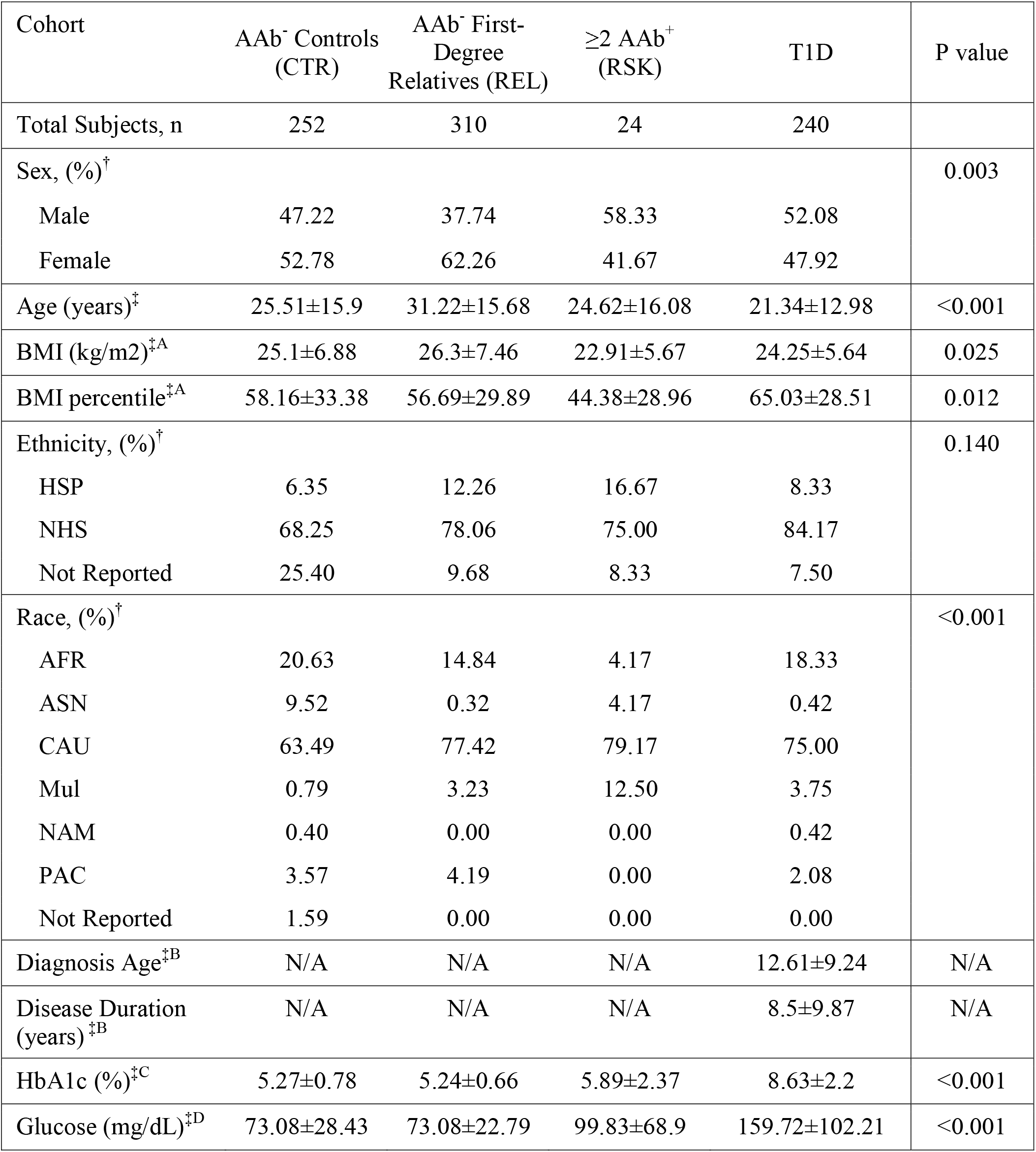
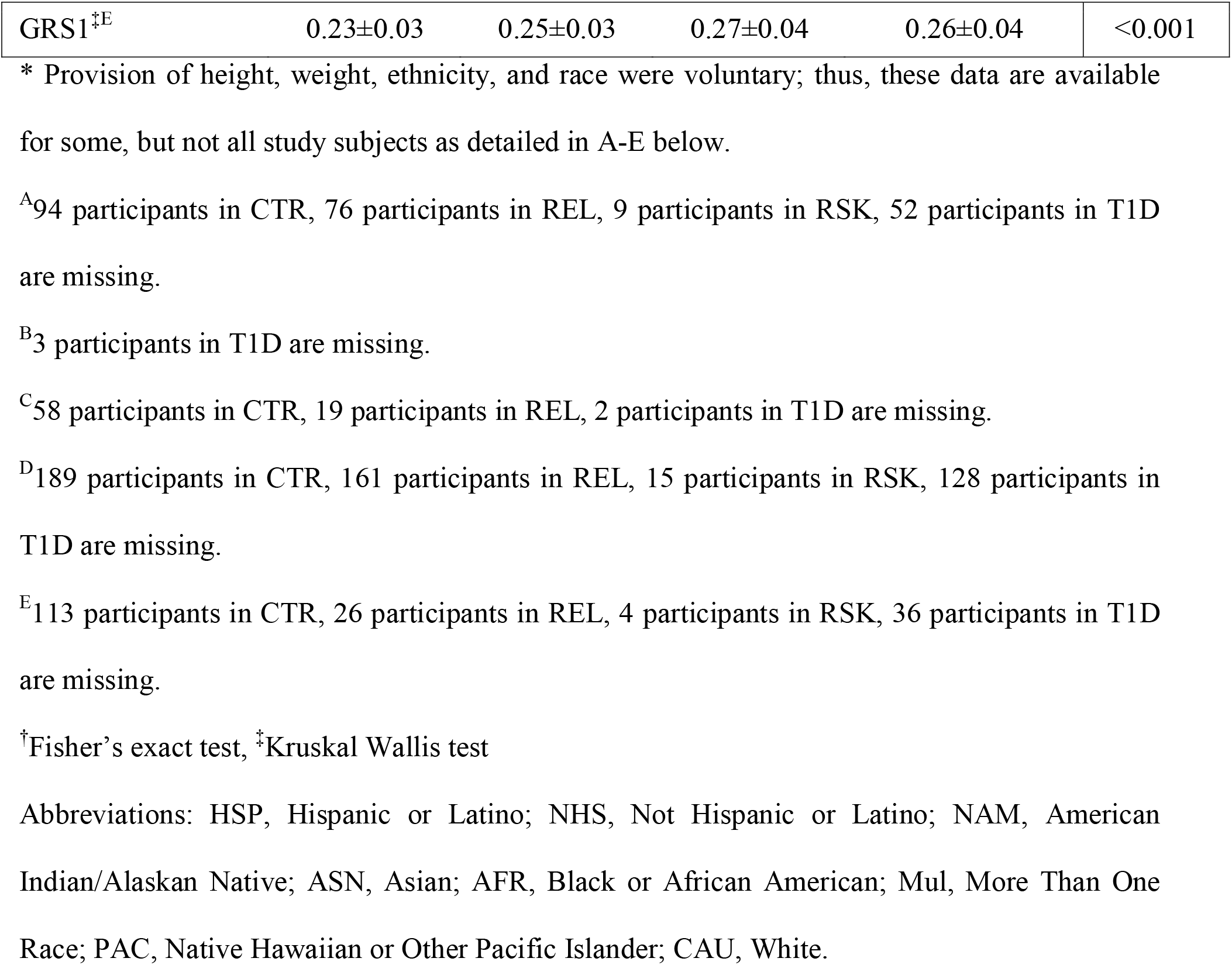
Demographic and clinical information. Data are presented as percentages (%) or mean ± SD.

### Altered age-associated immune trajectories in T1D

To assay the impact of age on the composition of the peripheral immune system, we characterized major trajectory patterns of the 172 flow cytometry outcome measures in healthy, unaffected islet AAb-CTR and REL between 5-75 years of age. Smoothing splines were fit to model each phenotype’s trajectory over age. Hierarchical clustering analysis of normalized, centered and scaled phenotype trajectories (**Figure 2A**) revealed four distinct patterns: 1) increasing linear (66 phenotypes), 2) upward parabolic (20 phenotypes), 3) decreasing linear (74 phenotypes), and 4) stable (12 phenotypes) relationships with age (**Figure 2B**). Using these defined cluster assignments (**Figure S7**), we also fit smooth trajectories on T1D samples and overlaid these with the splines for AAb-CTR and REL (**Figure 2C-D**). When organized using the clustering structure from the unaffected individuals, the T1D immune trajectories appeared to have similar overall trends (**Figure 2C**). Indeed, trajectory-specific difference revealed that the vast majority (86%) of phenotype trajectories initially trend in the same direction (increasing or decreasing), and those with differing initial trends cross at least twice, indicating sampling variation in the trajectory rather than disease-driven divergence. However, a direct comparison between T1D and AAb-trajectories revealed an average upward phenotypic shift of T1D in Cluster 1 (p<0.001) and a downward shift of T1D in Cluster 3 (p=0.034, **Figure 2D**). Over time, trajectories tended to shift further apart (**Figure S8**). Together, these observations suggest that individuals with T1D exhibit distinct age-dependent alterations in immune trajectories.

**Fig. 2.**
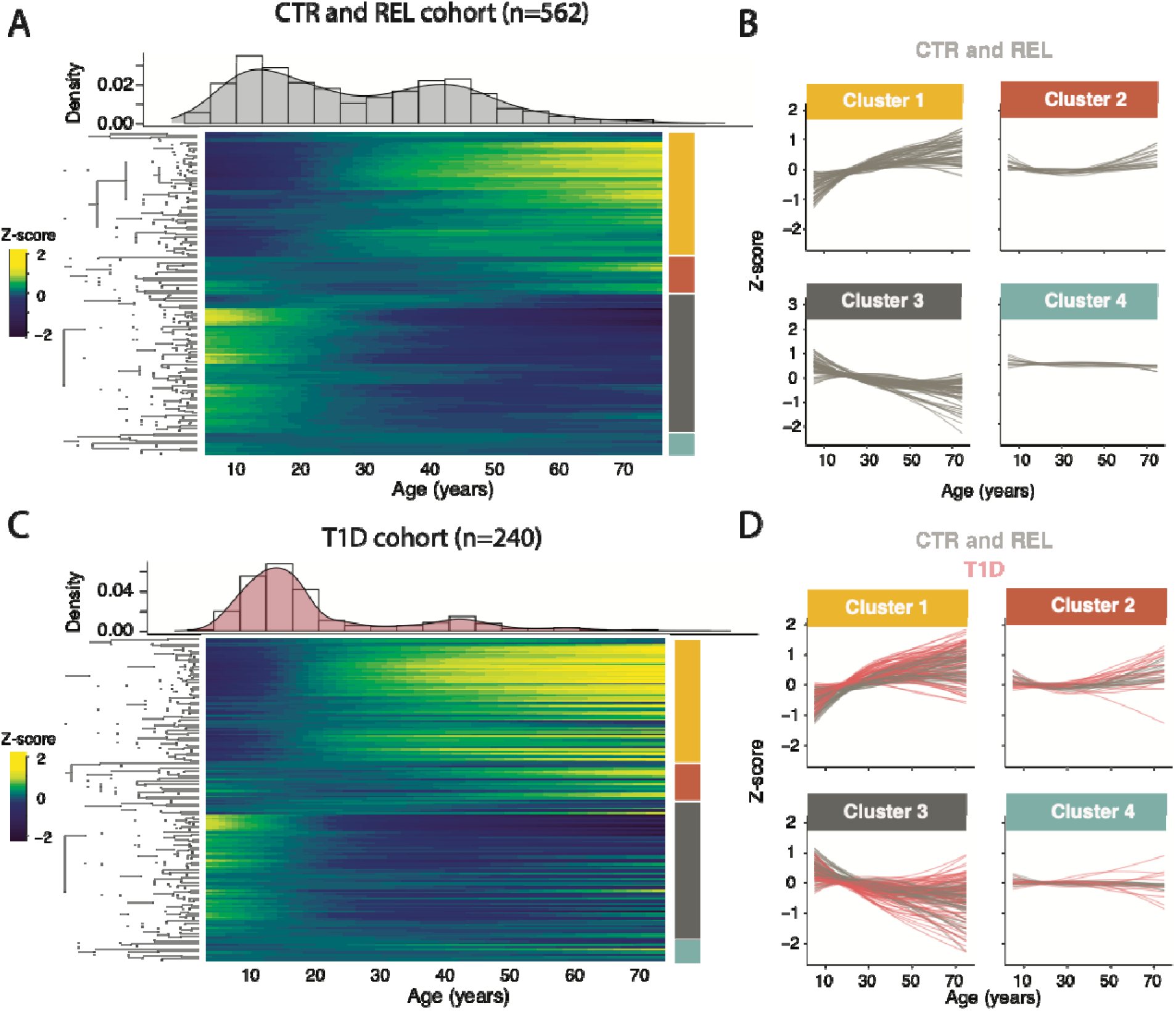
Altered immunophenotype trajectories in T1D. (**A**) Heatmap of smoothed phenotype trajectories as a function of age in AAb-individuals between the ages of 5 and 75 years. The age distribution of the cohort within this age range is shown as a histogram along the top axis. Immune cell phenotypes were clustered into four distinct groups (denoted by right axis colors) using hierarchical clustering (dendrogram shown on the left axis). (**B**) Line plots of each smoothed phenotype as a function of age demonstrate the distinct dynamic behavior within each of the four clusters. (**C**) Heatmap of smoothed phenotype trajectories as a function of age in T1D individuals with the rows in the exact order a indicated by the dendrogram in panel (A). (**D**) Line plots of each smoothed phenotype as a function of age as shown in panel (B) with the T1D smoothed phenotypes overlaid in red. (See also Figures S7-S8).

### Accelerated immune aging in T1D

To more specifically quantify the extent to which T1D immune profiles deviate from healthy individuals across chronologic age, we created a model to estimate an “immunological age” parameter from our CBC and flow cytometry data. Due to hierarchal dependence and correlation among flow cytometry readouts (**Figure S9**), we trained a random lasso (35) model on islet AAb-CTR to identify phenotypes predictive of age in individuals without autoimmunity. Our approach retained 69 immune features (**Figure 3A**) with a test set prediction performance of R^2^=0.70. Model consistency was observed by training on a combined AAb-cohort in which 45 variables were commonly retained and R^2^ = 0.70. Many of the phenotypes associated with age are consistent with the literature, providing support for our proposed immune age model. For example, B cell frequency decreased, with expected shifts from naïve to memory populations in both T and B cells over the lifespan (36–38). CD4^+^ T cells increased concomitant with a decline in CD8^+^ T cell frequency, consistent with the CD4/8 ratio increasing with age (39).

**Fig. 3.**
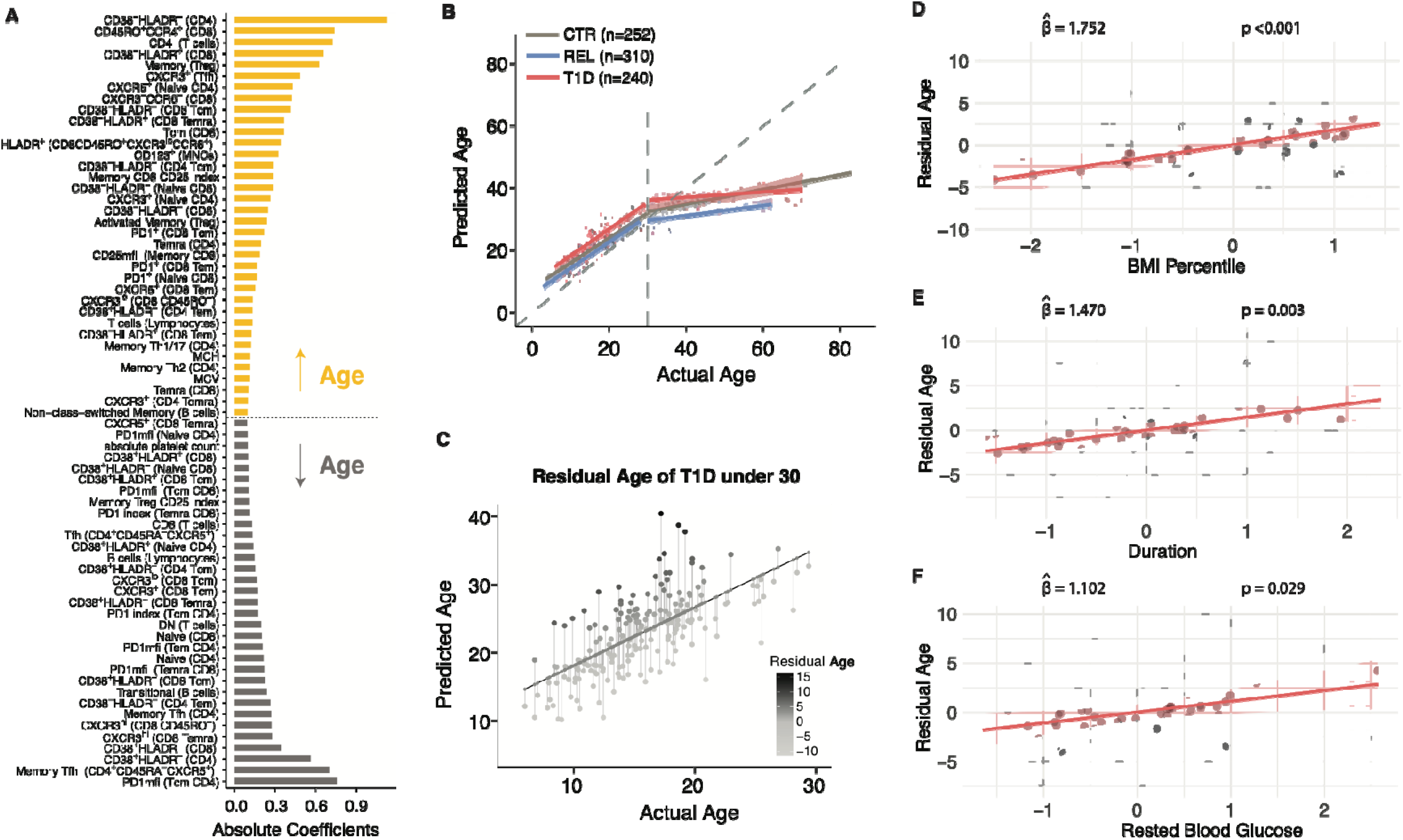
Immunophenotype age modeling reveals accelerated aging in T1D. (**A**) Averaged coefficients from the random lasso model are shown for all phenotypes above an empirically estimated threshold. The yellow phenotypes represent those increasing with age and gray denotes phenotypes decreasing with age. (**B**) The random lasso model was used to estimate an immunological predicted age. Predicted immunological age is shown for all CTR (gray), T1D (red), and AAb-REL (blue). The correspondence with the predicted age with chronological age is shown using a piece-wise regression model with a break at chronological age 30. (**C**) Residual immunological age is calculated the residual from a linear regression of predicted age and chronological age. (**D**) The partial regression plot between residual age and BMI percentile is shown for the multivariable regression model, along with the standardized coefficient and p-value. (**E**) Similar to panel D for T1D duration. (**F**) Similar to panel E for rested blood glucose. (See also Table S2 and Figures S10-S11).

We then applied our model to the full AAb-CTR cohort, along with the AAb-REL and T1D cohorts to examine differences in predicted age and chronological age. We observed the highest predictive performance of the immune age model among younger individuals regardless of risk cohort, which we confirmed by training two separate random lasso models for AAb-CTR younger than the age of 30 and those older (R^2^ = 0.66 versus R^2^ = 0.56). Given that the model plateaus after age 30 (**Figure 3B**), we focused the rest of our analysis on subjects under 30 years of age, where we observed the T1D group displayed accelerated immune aging relative to both AAb-CTR and REL, with an average increase of 3.36 years (p<0.001, **Figure 3B**). The observed accelerated immune aging in T1D did not appear to be related to history of CMV infection, which, perhaps surprisingly (24; 40), was not a significant driver of accelerated immune aging in our young cohort (average increase of 1.62 years associated with CMV infection history, p=0.086, **Figure S10**). Nor was the average increase of 3.36 years associated with demographic variables such as sex, race, or ethnicity when included in a multivariable regression model of predicted age. Clinical covariates BMI percentile, hemoglobin A1c (HbA1c), and rested blood glucose were significantly associated with accelerated immune aging in T1D (p<0.001, p=0.035, and p=0.007), perhaps not surprisingly since these values differed significantly across risk groups (**Table 1**). In addition to the age shift in T1D, we observed increased variation in the age predictions. We explored nine disease-relevant features for potential associations in the residual predicted age in T1D patients (**Figure 3C; Table S2**). While polygenic T1D risk (genetic risk score [GRS1] (5)), participant sex, self-reported race and ethnicity were not associated with residual age (**Table S2**), we found significant associations with disease duration, BMI percentile, and rested blood glucose (**Figure 3D-F**). Similar associations with BMI percentile and blood glucose were not observed in age residuals in AAb-CTR or REL (**Figure S11**). The standardized regression coefficients in the multivariable model indicated that BMI percentile had the largest contribution (β=1.75), followed by disease duration (β=1.47), and rested blood glucose level (β=1.10). The increased residual age variations observed in T1D compared to AAb-individuals and their association with clinical features reflects the additional immunological burdens associated with T1D disease.

### Age correction of immunophenotyping data in a T1D prediction model

Given the diverse and heterogeneous phenotypic distributions across the 192 flow and CBC measures, along with their non-linear association with age, we chose to use a semi-parametric modeling framework to investigate the specific flow cytometric readouts contributing to the accelerated immune aging observed in T1D. We obtained age-corrected phenotype data using generalized additive models for location, shape, and scale (GAMLSS) (41; 42). Using this approach, we modeled each phenotype as a smooth-function of age using the combined AAb-CTR and REL cohorts, obtained the fitted distribution parameters for all individuals ages, which we then used to obtain the centiles of their phenotypes from the estimated cumulative distribution function. The advantage of this approach is that in addition to allowing for age-adjusted testing of phenotypes between risk groups, it also provides the framework for building an immunophenotype centile reference range. We built an R/Shiny user interface (ImmScape; https://ufdiabetes.shinyapps.io/ImmScape/) for interactive exploration and visualization of raw data and age-adjusted centiles generated within this study, as illustrated in **Figure 4A-C**. With these age-corrected data, we were able to identify 26 features that were significantly increased and 22 that were significantly decreased in T1D as compared to CTR subjects following multiple testing correction (**Figure 4D, Table S3**). Significantly modulated phenotypes were present from all six flow cytometry panels and the CBC data (**Figure S9, Table S1**). To quantify the usefulness of the age-adjustment in identifying disease relevant phenotypes, we used logistic regression to predict disease status (T1D versus AAb-CTR) using the 48 T1D associated phenotypes and obtained an area under the receiver operating characteristic curve (AUROC) of 82.3%. This is an improvement over similarly constructed prediction models on all age-corrected phenotypes (AUROC=79.6%) and on all uncorrected phenotypes (AUROC=76.8%).

**Fig. 4.**
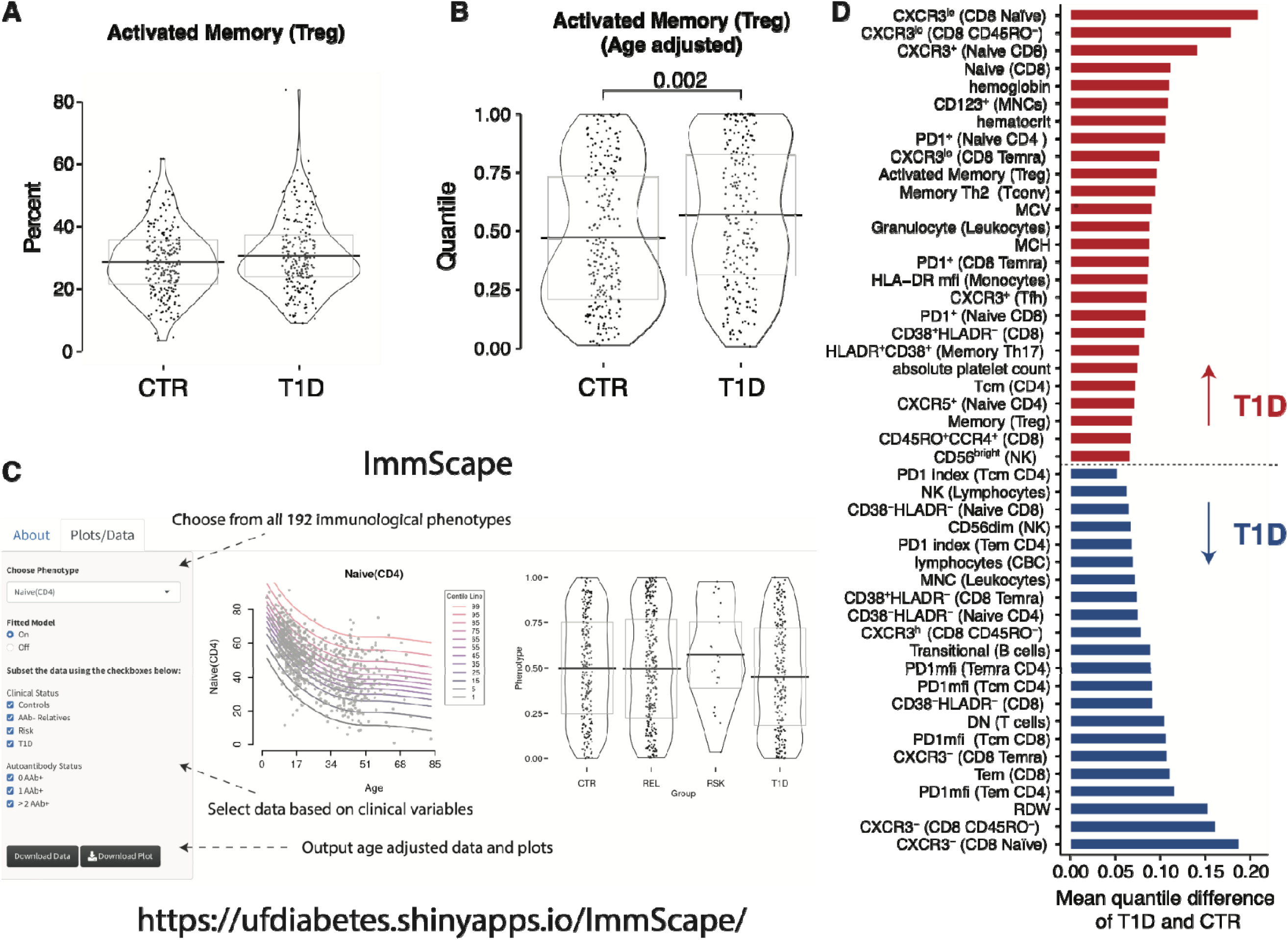
Age corrected phenotypes reveal T1D specific differences. As an example of the utility of our model, (**A**) in the uncorrected data, there is no significant difference detected between T1D and CTR. (**B**) Using the GAMLSS corrected data, there is a significant difference between T1D and CTR. (**C**) All age-corrected data is available for download and analysis via the ImmScape R/Shiny application. (**D**) The age-corrected quantile values for T1D versus CTR individuals were compared using a non-parametric Kruskal Wallis test and post-hoc Dunn’s test with a Benjamini-Hochberg multiplicity adjustment. Phenotypes that are increased in T1D (regardless of age) versus CTR are shown in Red. Phenotypes with decreased values in T1D versus CTR are shown in Blue. (See also Table S3 and Figures S9-S11).

### Immune features influenced by age and T1D status

Altogether, 46 measured variables were significantly modulated with age alone versus 25 with T1D status, while 23 readouts had shared contributions from both age and T1D (**Figure 5**). Many of our initial observations from flow cytometry and CBC data confirm previously reported findings in T1D. For example, the proportion of naïve CD8^+^ T cells was increased, while the CD8^+^ T effector memory (Tem) cell population exhibited decreased frequency in T1D (43) (**Figure S12A-B**). Moreover, we observed reduced frequencies of CD8^+^ T cells lacking the activation markers CD38 and HLA-DR in T1D with concomitant increase in CD8^+^CD38^+^HLA-DR^-^ cells (44) (**Figure S12C-D**). We also observed a decrease in CD56^dim^ NK cells with expected concomitant increase in CD56^bright^ NK cells in T1D subjects, in agreement with two publications (44; 45), but in conflict with another (46), as well as a trend (p=0.095) in ≥2AAb+ RSK subjects (**Figure S12E**). Furthermore, T1D subjects showed reduced frequency of transitional B cells as compared to AAb-CTR and REL (**Figure S12F**) (47). As summarized in **Figure 5**, we found an association of increasing non-class-switched memory B cells with age, while transitional B cells were found to decline with both aging and T1D (48; 49). Finally, CBC parameters showed both age and T1D dependent shifts: MCV and MCH increased with age and T1D, while hemoglobin and hematocrit increased in T1D subjects, findings supported by existing literature (50). Along with providing support for the aforementioned findings, we identified several repeating themes pertinent to immune dysregulation in T1D, namely, altered expression of the Th1-associated chemokine receptor CXCR3 and coinhibitory receptor PD-1 on multiple T cell subsets, as well as increased expression of HLA-DR on monocytes (**Figure 5**), which we explored further using our novel age-correction approach as described below.

**Fig. 5.**
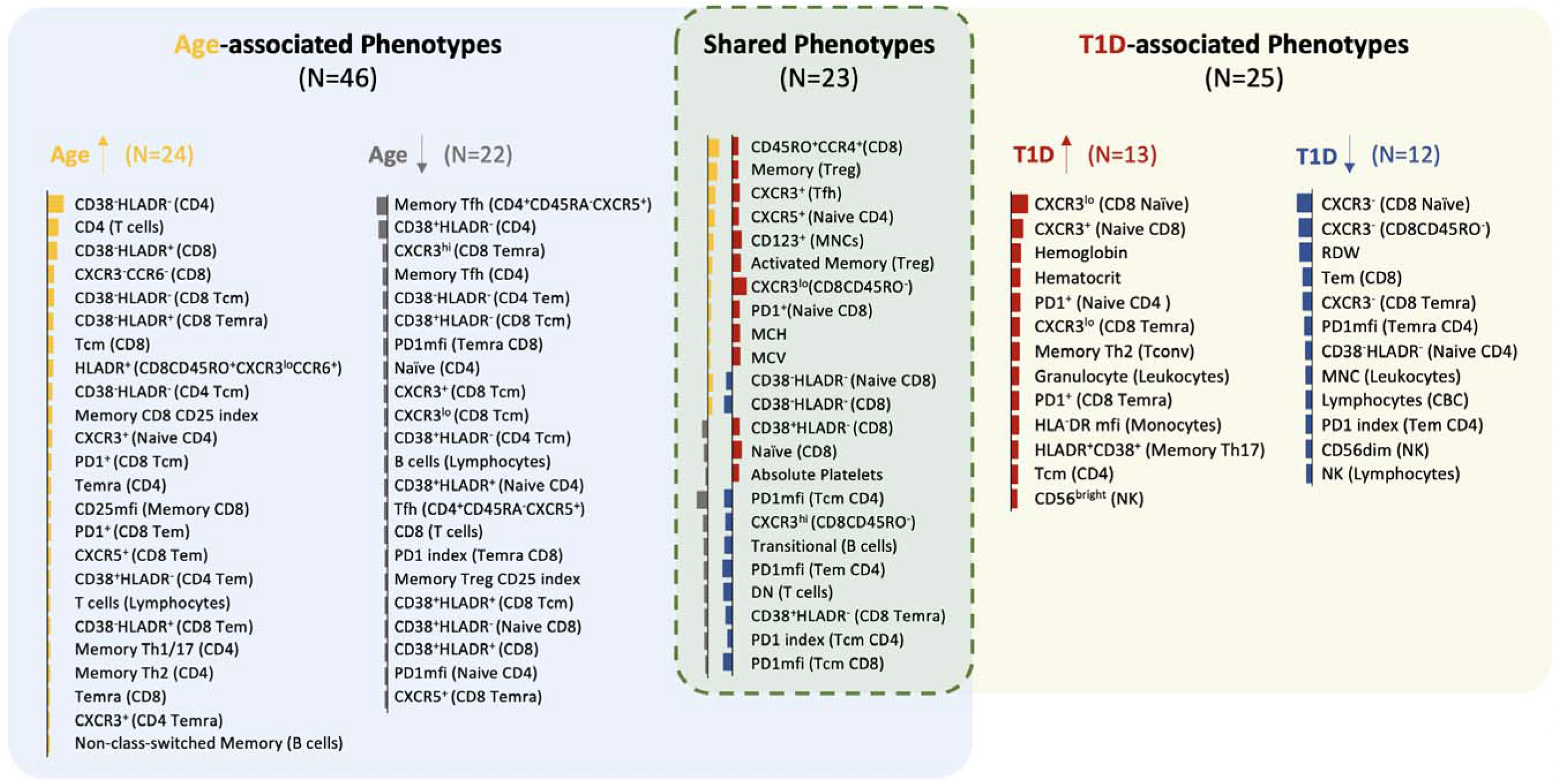
Overview of age- and T1D-associated phenotypes. A rectangular Venn diagram summarizes the phenotypes that are associated with age, T1D, or both. Phenotypes with unique association to age (left) or T1D (right) are shown in the non-overlapping areas, while the common phenotypes are displayed in the overlapping area (center). The total number of phenotypes that are “unique” or “common” to age or T1D are indicated in parenthesis. A color bar illustrating the magnitude and direction of effect on age or T1D is to the left of each phenotype (length of bar represents the effect size). The bar color indicates the phenotype is upregulated (yellow or red) or downregulated (gray or blue) in age or T1D, respectively.

### Increased CXCR3 expression on T cell subsets of T1D subjects

Of all parameters measured, the greatest mean difference between T1D and CTR (without a significant age association) was significantly increased frequency of CXCR3 expression among naïve CD8^+^ T cells (**Figure 5**), due to increased CXCR3^lo^ and decreased CXCR3^-^ subset percentages (**Figure 6A-B, Figure S13A-B**). CD8^+^ T effector memory CD45RA^+^ (Temra) exhibited the same patterns of increased CXCR3^lo^ and decreased CXCR3^-^ frequencies in T1D (**Figure 6C-D, Figure S13C-D**). Individuals with T1D also demonstrated elevated frequencies of CXCR3^+^ Tfh (**Figure 6E, Figure S13E**) as previously reported (44). Together, these data show a shift toward increased CXCR3 expressing populations across T cell subsets in T1D.

**Fig. 6.**
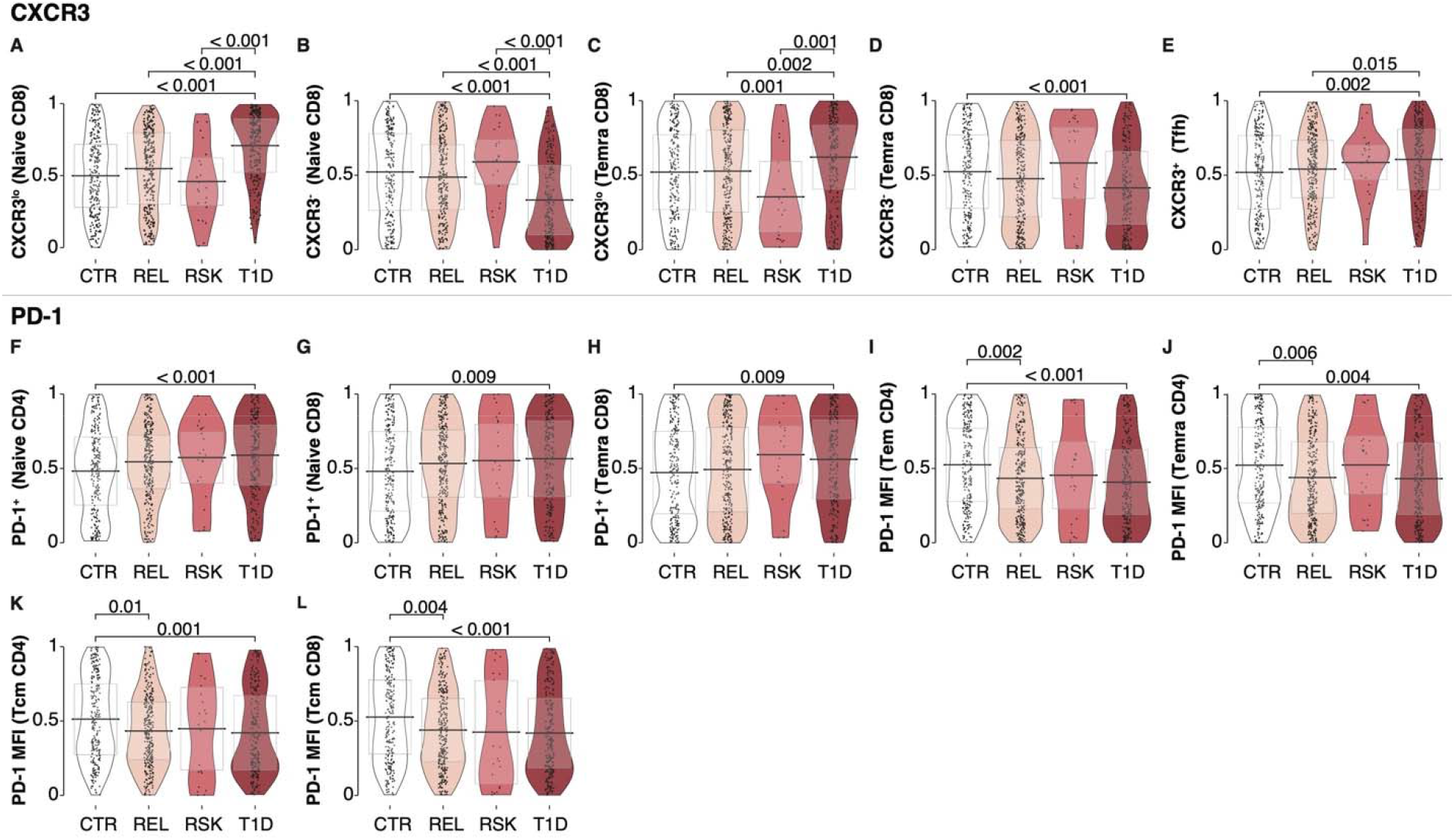
Increase in CXCR3-expressing T cell subsets and increased frequency – albeit lower intensity - of PD-1 expression in T1D. Age-corrected quantile values for (A-E) CXCR3^lo^, CXCR3^-^ or CXCR3^+^ frequency, (F-H) PD-1^+^ frequency, and (I-L) PD-1 MFI on T cell phenotypes. Significant p-values shown on graph. Testing was done using a Kruskal Wallis test with a post-hoc Dunn’s test and Benjamini-Hochberg multiplicity adjustment (See also Figure S13).

### Altered PD-1 expression on T cells from T1D subjects

The next major category of phenotypes involved altered expression of the coinhibitory receptor, PD-1, on T cell subsets (**Figure 5**). Despite observing significantly increased frequency of PD-1^+^ cells within naïve CD4, naïve CD8, and CD8^+^ Temra subsets from T1D subjects (**Figure 6F-H**, **Figure S13F-H**), PD-1 expression intensity (MFI) was decreased in T1D subjects on the majority of subsets analyzed: CD4^+^ Tem, CD4^+^ Temra, CD4^+^ Tcm, and CD8^+^ Tcm (**Figure 6I-L**, **Figure S13I-L**). Importantly, due to age-associated changes in PD-1 expression across all clinical groups (**Figure 3A**), a number of these T1D-associated differences were most apparent following age correction (**Figure 6G,** 6J-L**, Figure S13G, S13J-L**). Interestingly, PD-1 MFI was also significantly decreased on memory CD4^+^ and CD8^+^ T cell subsets of AAb-REL as compared to CTR (**Figure 6I-L**, **Figure S13I-L**), suggesting a potential genetic predisposition to altered expression of PD-1.

While some studies in small racially homogenous cohorts have associated single nucleotide polymorphisms (SNPs) in the *PDCD1* locus with T1D (51–54), large genome-wide association studies (GWAS) in European cohorts have failed to replicate these findings (31; 55). We surmised that a *PDCD1* SNP that has not been identified in GWAS efforts may be enriched in T1D, AAb+, and AAb-REL participants from our trans-ancestral cohort (**Table 1**). To test this hypothesis, we downloaded all *PDCD1* expression quantitative trait loci (eQTL) for whole blood from the Genotype-Tissue Expression project (GTEx) (56) and tested for increased presence of each SNP in T1D versus CTR groups via logistic regression. The top enriched SNP was rs6422701, with the T allele being overrepresented in the T1D cohort (**Table S4**). The T allele of rs6422701 was associated with decreased *PDCD1* mRNA expression in whole blood in GTeX (**Figure S14A**). Indeed, this translated to significantly decreased PD-1 expression on CD4 Tem and CD4 Temra, but not CD4 Tcm or CD8 Tcm, in CTR and AAb-REL with the T allele (**Figure S14B-E**), partially mirroring observations in REL vs CTR overall (**Figure 6I-L**, **Figure S13I-L)**. Together, these findings support the hypothesis that decreased PD-1 expression on memory T cells may contribute to T1D pathogenesis.

### Increased HLA-DR expression on monocytes with T1D-associated HLA-DR4 genotype

While most of the immune features associated with T1D (**Figure 5**) were comparable between AAb-CTR and AAb-REL (**Figure 3B**, **Figure 6A-H**), we noted that some phenotypes also had significant differences in these groups (**Figure 6I-L**). This led us to ask whether genetic loci contributing to risk for T1D, which are enriched in REL (5), might predispose individuals toward developing such immune subset perturbations. Thus, we performed QTL analysis on genetically unrelated subjects to test for associations between age-corrected flow cytometric phenotypes and genotypes at 240 T1D risk variants (29–31) found at ≥5% minor allele frequency (MAF) in our cohort, while considering RSK or T1D status, sex, and population stratification as possible covariates. Following adjustment for multiple testing of genotypes and phenotypes, a significant association (false discovery rate [FDR] < 0.05 corrected by Benjamini-Hochberg multiplicity adjustment for genotypes and phenotype) was observed between the rs7454108 T1D-risk genotype and increased HLA-DR MFI on monocytes (**Figure 7A, Table S5**). As rs7454108 tags the high-risk HLA-DR4-DQ8 haplotype (29; 57; 58), we asked whether other HLA haplotypes carrying strong risk or protection from T1D likewise impacted this phenotype. However, we did not find evidence of association with monocyte HLA-DR MFI for either the high-risk HLA-DR3-DQ2 haplotype (rs2187668 (5; 29)) or the dominant protective HLA-DR15-DQ6 haplotype (rs3129889 (5; 29)) (**Figure 7A**). Although HLA-DR expression levels on monocytes did not show age dependence (**Figure 7B**), we found that the GAMLSS-corrected data demonstrated evidence of a genotype-dosage effect (**Figure 7C**). Specifically, HLA-DR MFI was increased in subjects heterozygous for HLA-DR4 as compared to those carrying other HLA class II genotypes (DRX/X), and this was further increased in subjects homozygous for HLA-DR4 (**Figure 7C**). Notably, the association between HLA-DR4 genotype and HLA-DR MFI on monocytes was present in all groups across the spectrum of T1D development, suggesting that this genetic driver of immune phenotype may act independently of AAb positivity or disease status (**Figure 7D-G**).

**Fig. 7.**
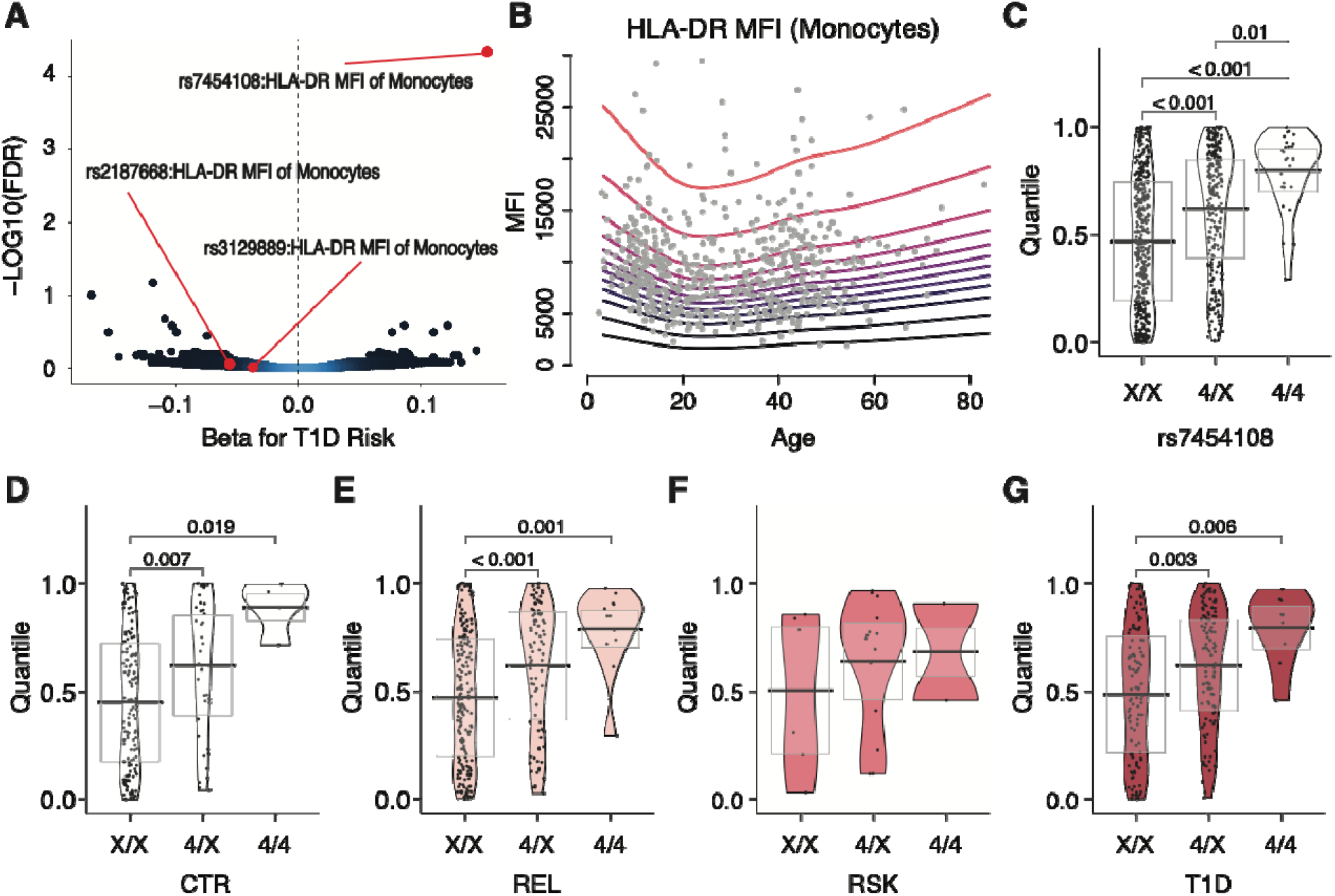
Increased HLA expression on monocytes in HLA-DR4 individuals. (A) Volcano plot showing quantitative trait locus (QTL) analysis results of all flow cytometry phenotype versus T1D risk loci. Associations shown according to direction and effect size (beta) of each single nucleotide polymorphism (SNP) on T1D risk. Blue designates higher and black designates lower data density. Associations between HLA-DR mean fluorescenc intensity (MFI) on monocytes and tag SNPs for HLA-DR4 (rs7454108), -DR3 (rs2187668), and -DR15 (rs3129889) T1D risk or protective class II HLA alleles highlighted in red. FDR = false discovery rate. (**B**) The GAMLSS model fit on all AAb-individuals (CTR and REL combined) in order to correct for age. Quantiles of HLA-DR MFI on monocytes in (**C**) whole cohort, (**D**) CTR, (**E**) AAb-REL, (**F**) ≥2 AAb+ RSK, (**G**) and T1D participants according to number of copies of HLA-DR4. X = any HLA-DR allele other than DR4. Significant p-values shown on graph. Testing was done using a Kruskal Wallis test with a post-hoc Dunn’s test and Benjamini-Hochberg multiplicity adjustment. (See also Table S5).

## DISCUSSION

In this study, we conducted flow cytometric analysis of peripheral blood from a well-characterized cross-sectional cohort (*n*=826), including healthy CTR, volunteers with T1D, as well as individuals at varying degrees of risk for T1D development, to understand how risk for autoimmune diabetes intersects with age to impact the broad immune landscape covering major subsets of the innate and adaptive immune system. We used these data, along with extensive genomic and clinical metadata, to visualize immune system changes across the human lifespan. This process revealed striking dynamics of immune age, primarily within the adaptive arm of the immune system and particularly in the first three decades of life, which could be used to predict chronological age with a high degree of accuracy in CTR. This resulting dataset corroborates a list of cellular subsets and phenotypic markers that change dramatically as a function of age (e.g., increased CD4^+^ T cells and decreased frequencies of B cells and CD8^+^ T cells, with shifts from naïve to memory populations in both T and B cells over the healthy human lifespan (36–39)). This complements prior descriptions of age-associated serological changes (59) and alterations in the epigenetic DNA methylation “clock” (60) that have been well characterized. Moreover, our data fill important gaps from prior studies (13; 61–63) to help understand changes in the pediatric immune system– a critical and dynamic time period for understanding normal frequencies and cellular distributions.

When our immune age algorithm was applied to subjects with T1D, the model revealed accelerated immune aging and numerous cellular immune phenotypes associated with T1D after GAMLSS age correction. Our age-correction model thus enabled the assessment of phenotypic changes as influenced by T1D, which is challenging in cross-sectional studies where age distributions of donors are often skewed. The analytical approaches described herein can be used to facilitate the inclusion of more subjects into study populations by enabling comparisons across poorly matched or unbalanced cohorts through the correction of age as a key covariate influencing the frequency and phenotype of major immune subsets. This is of particular value to biomarker studies in diseases impacting pediatric subjects, where sampling of healthy age-matched CTR is often limited. Beyond this, using our cross-sectional immunophenotyping dataset, we built a model capable of predicting T1D status with internal prediction performance at 82.3% accuracy. Evaluation of this model in larger pre-T1D cohorts and with longitudinal sampling will be necessary to validate its possible utility for T1D prediction and monitoring.

There are a number of reasons to consider age as a key aspect of T1D pathogenesis. Importantly, the first hallmark of β-cell autoimmunity (i.e., emergence of multiple islet AAb) often occurs within the first two years of life (6). A younger age-of-onset has also been associated with the highest risk HLA-DR3/4 diplotype (5), a prominent T and B cell insulitic lesion in the pancreas of T1D organ donors (9; 10), a bias toward IFN-γ:IL-10 T cell autoreactivity (64), and more acute clinical loss of endogenous C-peptide (29). While other studies have considered the impact of age on the changing immune system in T1D (65), we believe our study is unique by covering a large cohort, including a small group of rare ≥2AAb+ (i.e., pre-T1D (27)) participants, with subject ages spanning multiple decades, and implementing a strategy to account for age differences.

In an effort to explain the observed accelerated aging, we examined T1D GRS1 (5) and clinical covariates associated with disease duration and glycemic dysregulation. Modest associations with BMI percentile, disease duration, and rested blood glucose level were found, but polygenic risk did not contribute significantly to the observed change in immune age. Thus, we put forth that accelerated immune aging observed in T1D likely results from chronic inflammation, as has been described previously (66–70), and/or hyperglycemic stress. High-risk subjects who progressed to disease in The Environmental Determinants of Diabetes in the Young (TEDDY) cohort displayed chronic enteroviral shedding (71), evidence of persistent infection and consequent inflammation. In addition, Carey and colleagues found increased risk of infection in total diabetes participants when compared to CTR, but also in T1D when compared to type 2 diabetes, highlighting that individuals with T1D potentially suffer from multiple impacts to their overall immune function (72).

Within this study, we replicated multiple findings in smaller cohort studies of T1D subjects compared to CTR (43–45; 47–50), as well as immune aging studies (36–39), providing support for both the age prediction and T1D prediction capacity of the model generated herein. Moreover, we identified novel phenotypes of immune system changes with age and T1D, and have made this dataset readily accessible in an interactive format for public visualization and comparisons to other diseases and cohorts of interest. Phenotypic shifts associated with both aging and T1D generally reflected accelerated aging in their directionality. However, two age-associated phenotypes of note were reversed in subjects with T1D relative to aging trends: naïve CD8^+^ T cell and CD8^+^CD38^+^HLA-DR^-^ T cell frequencies were increased in T1D, despite decreasing with age in CTR. Notably, these phenotypes may reflect the same cell subset, considering that the majority of naïve CD8^+^ T cells are CD38^+^HLA-DR^-^ (**Figure S3**). In the periphery, in the absence of infection or inflammation, naïve T cells are largely maintained through homeostatic expansion and tonic T cell receptor (TCR) signaling (73). CD8^+^ T cells, despite exhibiting evidence of more frequent clonal expansion, usually represent a shrinking proportion of the T cell pool over time, perhaps due to post-expansion contraction and cell death (74). Their increase in the periphery in T1D subjects could be explained by several potential mechanisms, including increased thymic output, increased expansion, and/or differentiation to stem cell memory T cells (75), which could not be distinguished from naïve T cells in the panels used herein.

The observed increase in CXCR3 expression across multiple T cell subtypes could influence maturation potential, trafficking, and/or polarization to promote disease pathogenesis. Findings from murine models corroborate a role for CXCR3 in T cell trafficking to the islets and autoimmune diabetes development (76), indicating this Th1-skewing as a globally dysregulated phenotype in T1D. Milicic *et al.* similarly reported higher CXCR3^+^ memory T cell frequencies in high-risk AAb+ REL as compared to low-risk relatives (77), suggesting linkage to disease processes (77). CXCR3^+^ naïve T cells have been previously described in mice, are reported to be enriched in autoreactivity, and are thought to contain higher-affinity (CD5^hi^) autoreactive TCRs (78). Additionally, expression of CXCR3 on naïve CD8^+^ T cells has previously been associated with enhanced potential to differentiate into an effector phenotype (79; 80) along with enhanced capacity for IL-2 and TNF production post-stimulation (79). Confirmatory markers for stem cell memory T cells (CD45RA^+^CXCR3^+^CCR7^+^**CD27^+^CD28^+^CD95**^+^) (75) were not included in this analysis, and reflect an active avenue of future investigation.

The observed shifts in PD-1 expression in T1D extend upon a previous study demonstrating decreased *PDCD1* mRNA and PD-1 protein levels in CD4^+^ T cells of T1D subjects (81) by defining the particular subsets on which PD-1 expression is downregulated (CD4^+^ Tem, CD4^+^ Temra, CD4^+^ Tcm, and CD8^+^ Tcm). These data provide further support for the notion that the PD-1 pathway may serve as a critical negative checkpoint for maintaining tolerance to islet β-cells (82). Moreover, our observation of reduced PD-1 MFI on these T cell subsets in REL suggest a potential genetic predisposition toward impaired PD-1 expression. Our finding regarding monocyte HLA-DR overexpression in HLA-DR4 subjects further highlights how a major risk locus can influence surface protein expression levels and extends upon the prior observation of this association in cord blood samples (83). Increased activation and/or antigen presentation afforded by the increased expression could theoretically shift the overall functional avidity of autoreactive TCRs and alter aspects of selection and cellular differentiation under inflammatory skewing conditions in T1D. These phenotypic observations regarding expression of CXCR3, PD-1, and HLA-DR warrant future targeted studies on the mechanism and downstream impact of the observed altered expression levels and implications for cellular function.

In T1D, we observed amplification of immune aging trajectories, but do not have sufficient data in very young subjects (e.g., birth to age seven) to understand when this trend is initiated, and did not assess longitudinal samples to compare to preclinical status. As result of this data sparsity at either end of the lifespan, our interpretation of findings applies most directly to the impact of diagnosed disease on immunophenotypes after age-correction of data. Association of immune phenotypes with progression of T1D in at-risk individuals (in this study, ≥2AAb+), would be best studied in longitudinal samples within individuals. Potential covariates of interest to immunophenotype shifts include pubertal status and time of blood sample draw (84), which were not recorded or included in the analysis. The breadth of this study is limited to the immunophenotypes and effector molecules investigated herein. Other immunological compartments of potential interest not explored in this study include mast cells, eosinophils, and basophils (85; 86), along with tissue resident populations (11; 87–89). Experiments to address these questions are ongoing.

Our efforts to identify a signature of immune system age, as well as cellular phenotypes associated with T1D, provide a number of additional targets for consideration in precision medicine-based therapies (e.g., CXCR3^+^ T cells, the PD-1 costimulatory axis, antigen presentation on monocytes, specifically in individuals with HLA-DR4). Moreover, these results provide a number of cellular and pathway targets for consideration in mechanistic studies of prior trials where efficacy differed according to the age of the trial participants (e.g., rituximab [anti-CD20, TN05], low-dose anti-thymocyte globulin [ATG, TN19], abatacept [CTLA4-Ig, TN09 and TN18]) (90; 91) and in those planned for future intervention studies. Importantly, immune-directed therapeutic interventions aimed at interrupting the autoimmune destruction of β-cells, at or prior to clinical diagnosis of T1D, often have outcomes impacted by the age of subjects at the time of drug treatment (reviewed in (92)). We suggest this observation reflects altered immune system constituents that shape therapeutic response (or lack thereof), depending on the target and drug mechanism-of-action. The approach of considering immune age, phenotype, and chronological age of trial participants may improve clinical response profiles and progress toward precision medicine-based strategies to prevent and reverse T1D.

## MATERIALS AND METHODS

### Study Design

Individuals with T1D, their REL, and CTR participants were recruited from the general population and outpatient endocrinology clinics at the University of Florida (UF; Gainesville, FL), Nemours Children’s Hospital (Orlando, FL), and Emory University (Atlanta, GA). Following procurement of written informed consent, peripheral blood samples were collected into the UF Diabetes Institute (UFDI) Study Bank from 826 non-fasted participants by venipuncture. Samples were collected in ethylenediaminetetraacetic acid (EDTA)-coated vacutainer tubes for flow cytometry, complete blood count, HbA1c, TCRβ sequencing, and genotyping assays; serum separator vacutainer tubes for islet autoantibody and CMV IgG antibody measurement; and sodium fluoride/potassium oxalate-coated tubes (BD Biosciences) for rested blood glucose quantification, and stored in accordance with institutional review board (IRB)-approved protocols at each participating institution. Samples were shipped or rested overnight prior to flow cytometric staining in order to standardize duration between collection and evaluation at UF (33). Data were collected from all incoming blood samples to the UFDI from 2014-2018, hence the lack of age-matching between clinical subgroups based on clinical status. Importantly, participants were generally healthy with no reported infection or malignancy at time of blood draw, and sample collection preceded the coronavirus disease 2019 (COVID-19) pandemic. Deidentified demographic and clinical information are presented in **Table 1**.

### Autoantibody Measurement

Islet Autoantibody Standardization Program (IASP)-evaluated enzyme-linked immunosorbent assays (ELISA) (93) were performed on serum to measure T1D-associated AAb reactive to glutamic acid decarboxylase 65 (GADA), insulinoma-associated protein-2 (IA-2A), and zinc transporter-8 (ZnT8A), which respectively performed with AUROCs of 0.936, 0.876, and 0.917 in the most recent IASP workshop. REL and CTR participants were considered at-risk for T1D (RSK) if possessing reactivity to at least two of the screened AAb specificities (27).

### Flow Cytometry

Rested whole blood samples were stained with six flow cytometry panels, five of which were adapted from the standardized HIPC recommendations (13) plus a Tfh-specific panel, as we have previously described (18; 32). Briefly, 200 µL of whole blood was stained with each panel of anti-human antibodies (**Table S6**) for 30 minutes at 23°C in the dark. Red blood cells (RBC) were lysed for five minutes at 23°C using 1-step Fix/Lyse Solution (eBioscience) and removed from the leukocyte suspension with three washes of staining buffer. Data were acquired on an LSRFortessa (BD Biosciences) within 24 hours of staining. Analyses were performed in FlowJo software (v9 and v10; BD Life Sciences) according to the gating strategies in **Figure S1-S6** to derive percentage of parent and geometric mean fluorescence intensity (gMFI) data.

### Complete Blood Count, HbA1c, and Blood Glucose Measurement

Rested whole blood samples were characterized using the Coulter Ac•T 5diff CP (Cap Pierce) Hematology Analyzer. Complete blood count (CBC) measurements included absolute count of red blood cells (RBC) and white blood cells (WBC), with absolute count and percentage of neutrophils, lymphocytes, monocytes, eosinophils, and basophils reported. Additional RBC parameters measured included hemoglobin (HGB) concentration, hematocrit (HCT), mean corpuscular volume (MCV), mean corpuscular hemoglobulin (MCH), mean cell hemoglobin concentration (MCHC), and red blood cell distribution width (RDW). Absolute count of platelets (PLT) and mean platelet volume (MPV) were also included. HbA1c was measured with the DCA Vantage Analyzer (Siemens) and rested blood glucose with the Contour Next EZ glucometer (Bayer).

### CMV Status

A random subset of the flow cytometry cohort (40.56% of the total cohort) was assayed for evidence of prior or primary CMV exposure by predicting serostatus from *TRBV* (TCRβ) sequences (Adaptive Biotechnologies) derived from peripheral blood mononuclear cells (PBMCs) (94; 95). CMV status was inferred using methods and trained model described by Emerson *et al.* (96). Upon testing, predictions over 0.5 were deemed CMV positive. Serostatus from a CMV IgG antibody ELISA (Zeus Scientific) was available for 71.04% of all samples that were TCR sequenced from our cohort. We noted reliable concordance between the two methods of CMV status classification, as evidenced by AUROC of 0.886. In cases of discrepancy (14.83% of subjects with data from both assays), CMV classification from ELISA superseded the TCRβ data as the result used for further analysis.

### Genotyping of T1D Risk Loci and Quantitative Trait Loci Analysis

Samples were genotyped at >985,971 unique loci using our custom single nucleotide polymorphism (SNP) array termed the UF Diabetes Institute UFDIchip (97), which includes the Axiom^TM^ Precision Medicine Research Array (ThermoFisher Scientific) plus all content from the ImmunoChip.v2.0 (98) as well as previously reported credible T1D risk variants (55). All quality control (QC) steps and QTL assessment were performed with plink 1.9 . T1D GRS was calculated as previously established (97).

QC measures were performed prior to association testing, as previously described (99). Subjects were excluded from analysis if any of the following conditions applied: >2% of directly genotyped SNPs were missing, genetically-imputed sex did not match reported sex, or heterozygosity rate differed ±3 standard deviations (SD) from the mean of all samples (2.91% of subjects excluded based on these criteria). Additionally, identity by descent (IBD) calculations were used to remove related individuals with pi-hat >0.2, randomly retaining one subject in each related pair (100), resulting in the exclusion of 40.56% of subjects. Genotypes at 277 previously curated T1D risk loci (29–31) were pulled directly from the UFDIchip or if missing from the chip, obtained from imputation to 1000 Genomes Phase 3 (v5) or Human Reference Consortium (vr1.1) using the Michigan Imputation Server (101), with imputed genotypes from the reference cohort providing the higher imputation quality used for QTL analysis (mean R^2^ = 0.952). SNPs that were missing in >2% of subjects, with a minor allele frequency (MAF) <5%, or failing Hardy-Weinberg equilibrium (HWE) at p<1e-6 were excluded. A linear regression analysis was performed using plink (102) for genotype association with GAMLSS-corrected flow cytometric phenotypes, with T1D status (AAb-with no family history of T1D [CTR], AAb-REL, ≥2 AAb+ RSK, T1D) and sex included as categorical covariates and ten multidimensional scaling (MDS) components as continuous covariates to account for population stratification. P-values were adjusted for multiple testing of genotypes and phenotypes to generate a FDR using the Benjamini-Hochberg method with R/p.adjust. A volcano plot was created using GraphPad Prism v7.0 depicting -log10(FDR) and the magnitude of the association between the phenotype and the T1D risk allele (beta coefficient) (29–31).

### Statistical Analyses

#### Phenotype dynamics and trajectories

Data are presented as mean ± SD, and all tests were two-sided unless otherwise specified. Technical and biological coefficient of variation (CV) in the flow cytometric assay were assessed on a cohort of 12 samples that were run in technical duplicates using GraphPad Prism software version 7.0. Spearman correlations between immune phenotypes and age were computed using R/pspearman.

For characterizing the flow cytometric phenotype dynamics over age in CTR and AAb-REL, missing phenotype data due to failure of visual QC validation of staining (<5.03% of data per phenotype) were median-imputed. Smoothing splines of each phenotype versus age were fit with three degrees of freedom using the smooth.spline R function (103).

To model the dynamics of each phenotype across age, we log transformed the imputed data, adding a constant equal to one or, for phenotypes with possible values less than 1, we added an additional constant to shift all values larger or equal to zero. We then z-transform scaled each phenotype prior to fitting a smoothing spline, as described above, across age with three degrees of freedom. The smoothed trajectory predictions were restricted to an age range between 5 and 75 years to avoid predicting outside the observed age range of our other cohorts present. The phenotype trajectories were then clustered using hierarchical clustering with the complete method and the Canberra distance metric to group phenotypes with similar trajectory patterns over age. Heatmaps of processed data from AAb-CTR and REL samples, ordered by subject age on the x-axis and dendrogram clustering on the y-axis, were created using R/gplots. R/ggplot2 and R/ggpubr were used to create figures of overlaid smooth splines of phenotypes for four clusters. The same procedures were applied to the T1D individuals to obtain T1D phenotype trajectories, and the T1D smoothed trajectories were plotted keeping the same y-axis order for comparison with age-related trajectories in AAb-CTR and REL.

### Comparing smooth trajectory fits

To compare the spline fits for the two cohorts (AAb-CTR and REL combined vs T1D), we computed the trajectory shift as the difference in the overall average trajectory value between cohorts. A t-test was used to test for significant shifts for each of the four clusters. To determine the initial trajectory direction, for each phenotype in each cohort we calculated the mean value of successive differences in the trajectory over years 5-15, so that a positive value indicated an increasing trajectory and negative value a decreasing trajectory. We estimated the number of times the trajectories crossed by examining the number of sign changes along the successive differences across the entire age range obtained.

### Immunophenotype age model

We used the random lasso (35) method on the CTR cohort to identify phenotypes associated with immune aging. We first imputed missing values within each phenotype to its median and then z-transform scaled each phenotype. With biological age as the response and all phenotypes as predictors, we repeated our random lasso procedure on 1,000 random train-test subsets with 80% of the individuals in a training set and the remaining 20% as a held-out test set. The first stage of the random lasso consisted of 1,000 bootstrap samples of the training set, with each bootstrap estimating the coefficients on 15-20% randomly selected phenotypes. The initial variable importance score for all phenotypes was calculated as the average coefficient value. In the second stage of the random lasso, another 1,000 bootstrap samples were taken from the training set and 10% of the phenotypes were fit in a lasso model using the initial importance score as the selection probability. For each data split, the overall variable importance score is obtained by averaging the bootstrapped coefficients. We obtained a final variable importance score by averaging over all 1,000 random data splits.

Of the 192 phenotypes evaluated, 182 had non-zero average coefficients from the random lasso procedure; thus, we further implemented a procedure to obtain an importance score cutoff. All variables higher than our cutoff threshold were considered predictive of age. To determine the optimal cutoff value, we ran 5-fold cross validation and evaluated the root mean squared error (RMSE) for each cutoff in the random lasso procedure. We estimated an elbow point per fold to minimize RMSE. We repeated this 5-fold cross validation 20 times, and the mean of elbow points was the final cutoff. The final linear model is the averaged coefficients from the random lasso models, to which we then applied to the AAb-CTR, AAb-REL, and T1D cohorts to obtain the predicted immunological age.

To internally validate our age prediction model, we similarly trained a random lasso model on additional cohorts: 1) the combined cohort of AAb-CTR and REL, 2) the AAb-CTR for individuals younger than 30, and 3) the AAb-CTR for individuals older than 30.

### Analysis of age prediction within T1D and control subjects

A linear regression model of predicted age was fit on the chronological age and a factor for clinical group (T1D versus all AAb-individuals). We then fit a multivariable model of predicted age on chronological age, sex, ethnicity, reported race, BMI percentile, HbA1c, rested blood glucose, and GRS. BMI percentile was calculated as previously described (104). Continuous variables were z-transform scaled to obtain standardized coefficients. To ensure a sufficient number of individuals were represented across risk groups, the model was limited to subjects with reported race of African American or Caucasian (see **Table 1** for sample sizes).

Within the T1D cohort, we again fit a multivariable model for the variables described above in addition to disease duration and diagnosis age. Due to the correspondence in our cohort between diagnosis age and age at sampling time, we used residual age as the outcome variable. We calculated the residual age as the residuals from a linear regression of predicted age and chronological age. This was done rather than calculating a delta age due to some remaining age association in the predicted age, and to more intuitively explain factors that associate with over/under-estimates of age within T1D. The standardized multivariable model was also fit to the AAb-CTR and AAb-REL cohorts separately.

### Adjusting for age using a GAMLSS model

In order to obtain age-adjusted centiles of immunophenotypes in CTR and T1D cohorts, we employed a weighted GAMLSS (41; 42). Using the CTR and AAb-REL, each phenotype was modeled as a cubic spline function of age using either a Box-Cox-t distribution or Normal distribution, depending upon the skewness of the phenotype distribution. A skewness cutoff of 0.5 was determined empirically, with distributions having skew larger than 0.5 fit using the Box-Cox-t (BCT) distribution. For phenotypes using the BCT distribution, values were shifted by a constant value to ensure positivity as necessary. Weights were assigned according to the age distribution density such that ages younger than 10 years were assigned a relative weight of 10, ages older than 70 years were assigned a relative weight of 0.1, with all other ages assigned a weight of 1. We then used the distribution function (pBCT or pNO) and the predicted parameter values from the weighted GAMLSS model to obtain age-corrected quantile values for all individuals. We note two exceptions to the above: 1) Memory Treg CD25 Index had a skewness of 0.54 but was better fit using the Normal distribution, and 2) CXCR3^hi^ (CD8 Temra) was fit using the unweighted GAMLSS model (all weights equal to 1) due to errors using the weighted model.

### Identifying T1D associated immunophenotypes

The age-corrected quantile values were used to compare T1D versus RSK, AAb-FDR, and unrelated AAb-CTR individuals. A non-parametric Kruskal Wallis test was performed for each phenotype followed by a post-hoc Dunn’s test with a Benjamini-Hochberg multiplicity adjustment. Multiplicity-adjusted P-values < 0.05 were considered significant.

### Immunophenotype prediction model

We first imputed missing values within each phenotype to its median and then z-transform scaled each phenotype. To handle the correlations and dependencies in the phenotypes, we used principal component analysis. The first 30 components were selected based on an elbow plot of explained variance and used as predictors in generalized linear model of T1D versus CTR status. The AUROC was averaged over 1,000 independent samplings of an 80:20 train-test set split.

## Supplementary Materials

Table S1. Proportions of CBC data falling outside of established 5^th^-95^th^ percentile reference ranges.

Table S2. Multivariable model of disease-relevant features with residual age in T1D individuals.

Table S3. Cellular subset frequencies and phenotypes assessed in peripheral blood of T1D and controls

Table S4. Logistic regression of Genotype-Tissue Expression project (GTeX) *PDCD1* expression quantitative trait loci (eQTL) in CTR versus T1D subjects.

Table S5. QTL analysis.

Table S6. Reagents and resources used in the study.

Figure S1. B cell panel gating.

Figure S2. Innate immune cell panel gating.

Figure S3. T cell panel gating.

Figure S4. Effector T cell (Teff) panel gating.

Figure S5. T follicular helper cell (Tfh) panel gating.

Figure S6. Regulatory T cell (Treg) panel gating.

Figure S7. Dendrogram of phenotype age trajectories.

Figure S8. Age-wise trajectory differences.

Figure S9. Correlations among significant age-corrected phenotypes.

Figure S10. Immune aging model with CMV status.

Figure S11. Residual age analysis in AAb-CTR and REL under 30 years of age.

Figure S12. Adaptive immune compartment findings confirming prior literature.

Figure S13. Raw data shows CXCR3 and PD-1 expression is increased on naïve T cells but decreased on memory T cell subsets in AAb-CTR, REL, ≥2 AAb+ and T1D individuals.

Figure S14. *PDCD1* expression quantitative trait locus (eQTL), rs6422701, associates with PD-1 mRNA and protein expression.

## Acknowledgments

The authors thank the sample donors and their families for providing blood samples, without which this work would not have been possible. Special thanks are extended to Kieran McGrail (University of Florida) for biorepository management, Jennifer Hosford (University of Florida) for participant recruitment, and Michael Clare-Salzler (University of Florida) for critical review of the manuscript. The authors thank the clinical staff at the University of Florida, Nemours Children’s Hospital, and Emory University for sample acquisition.

## Funding

National Institutes of Health grant P01 AI042288 (MAA)

National Institutes of Health grant R01 DK106191 (TMB)

National Institutes of Health HIRN CHIB grant UG3 DK122638 (TMB, CEM)

The Leona M. and Harry B. Helmsley Charitable Trust 2019PG-T1D011 (TMB)

JDRF Postdoctoral Fellowship 3-PDF-2022-1137-A-N (MRS)

Diabetes Research Connection project 45 (MRS)

JDRF Postdoctoral Fellowship 2-PDF-2016-207-A-N (DJP)

National Institutes of Health grant F30 DK128945 (PT)

National Institutes of Health grant T32 DK108736 (LDP)

National Institutes of Health grant F31 DK129004 (LDP)

## Author contributions

Conceptualization: DJP, DAS, MAA, TMB

Methodology: DJP, MRS, XD, MAB, RLB

Formal analysis: MRS, XD, RLB

Investigation: DJP, PT, JMM, LDP, KM

Resources: TMB, MAA, AM, LMJ, MJH, DAS

Data Curation: MRS, XD, DJP, RSM

Visualization: DJP, RLB, XD, MRS, PT

Funding acquisition: TMB, MAA, CEM, MRS, PT, LDP

Project administration: DJP, MRS, RLB, TMB

Supervision: DJP, RLB, MAB, TMB

Writing – original draft: MRS, MAB, XD, DJP, RLB

Writing – review & editing: MRS, XD, DJP, JMM, PT, ALP, LDP, KM, RSM, MAB, AM, PC, LMJ, CEM, CHW, MJH, DAS, MAA, RLB, TMB

## Data and materials availability

Data are available for visualization and analysis via an interactive R/Shiny application (ImmScape; https://ufdiabetes.shinyapps.io/ImmScape/) and from the corresponding author upon reasonable request. Flow cytometric data are in the process of curation and will be available at ImmPort (https://www.immport.org).

---

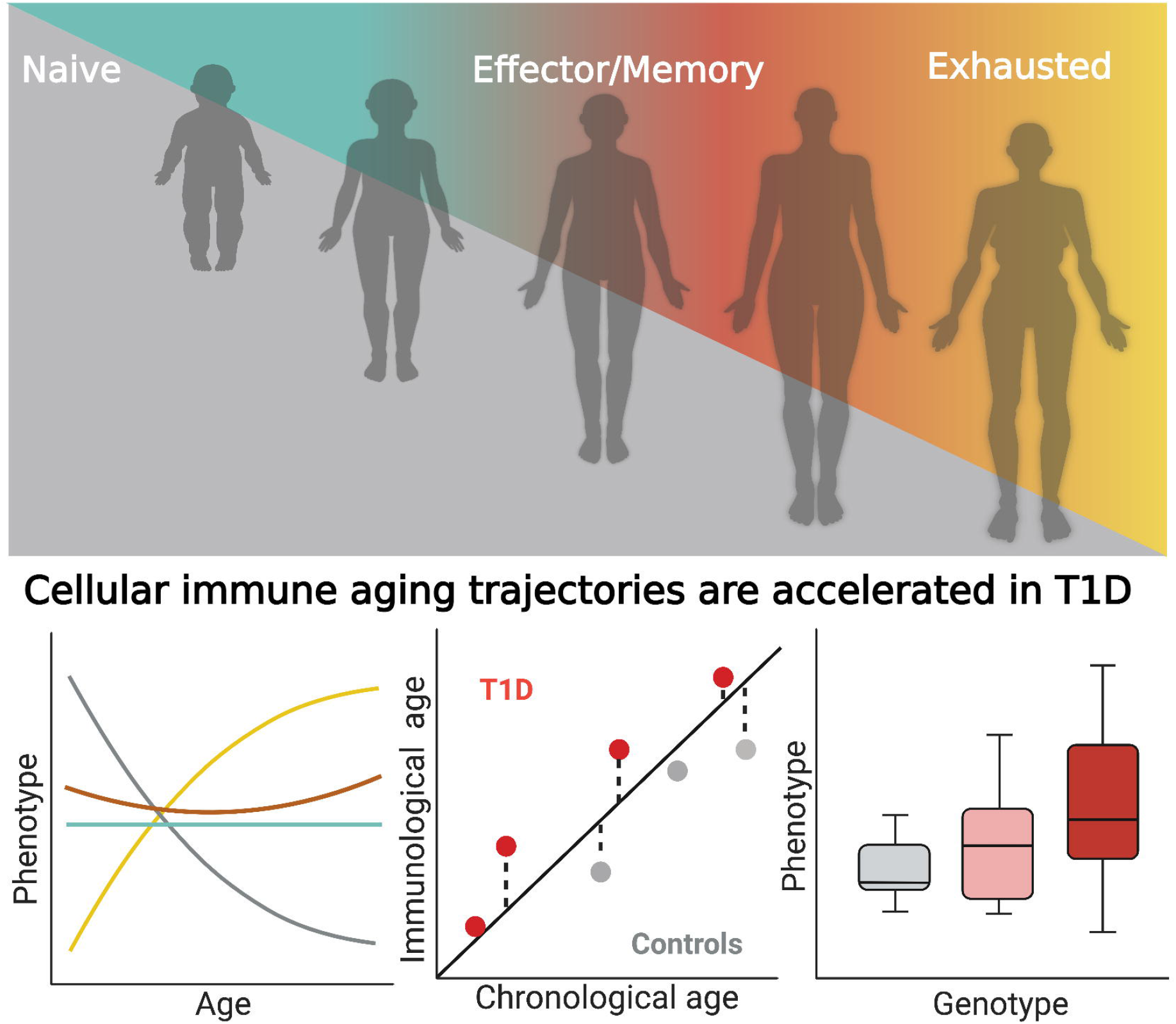

