## Supplementary Tables and Figures for "Human immune phenotyping reveals accelerated aging in type 1 diabetes"

**Supplementary Materials**

**Table S1. Proportions of CBC data falling outside of established 5^th^-95^th^ percentile reference ranges.** Percentage of autoantibody-negative (AAb-) control (CTR), AAb- first degree relative (REL), ≥2 AAb+ (RSK), and type 1 diabetes (T1D) participants with data below (Low, <5^th^ percentile), within (Normal, 5^th^-95^th^ percentile), and above (High, >95^th^ percentile) reference ranges reported for the instrument (Ac*T 5diff Cap Pierce Hematology Analyzer, Beckman Coulter). Data were analyzed by Chi-square tests with Bonferroni adjustment with significant differences, defined as adjusted p<0.05, shown in bold. (See also Figure 1)

| **CBC Phenotype** | **CTR (%)** | | | | **REL (%)** | | | | **RSK (%)** | | | | **T1D (%)** | | | |
| --- | --- | --- | --- | --- | --- | --- | --- | --- | --- | --- | --- | --- | --- | --- | --- | --- |
|  | **Low** | **Normal** | **High** | **Adj. p** | **Low** | **Normal** | **High** | **Adj. p** | **Low** | **Normal** | **High** | **Adj. p** | **Low** | **Normal** | **High** | **Adj. p** |
| WBC (10^3^/µL) | 6.2 | 87.1 | 6.7 | 1.000 | 2.4 | 89.7 | 7.9 | 0.950 | 4.5 | 90.9 | 4.5 | 1.000 | 4.2 | 89.7 | 6.1 | 1.000 |
| RBC (10^6^/µL) | 4.1 | 89.2 | 6.7 | 1.000 | 2.1 | 94.8 | 3.1 | 1.000 | 0.0 | 95.5 | 4.5 | 1.000 | 3.7 | 88.4 | 7.9 | 1.000 |
| HGB (g/dL) | 6.7 | 89.2 | 4.1 | 1.000 | 6.9 | 90.3 | 2.8 | 1.000 | 0.0 | 90.9 | 9.1 | 1.000 | 4.7 | 88.4 | 7.0 | 1.000 |
| HCT (%) | 6.2 | 86.1 | 7.7 | 1.000 | 4.5 | 85.9 | 9.7 | 0.105 | 0.0 | 81.8 | 18.2 | 0.930 | 3.3 | 84.2 | 12.6 | **<0.001** |
| MCV (fL) | 6.7 | 83.5 | 9.8 | 0.335 | 4.5 | 86.6 | 9.0 | 0.639 | 0.0 | 90.9 | 9.1 | 1.000 | 1.4 | 92.6 | 6.0 | 1.000 |
| MCH (pg) | 8.2 | 87.1 | 4.6 | 1.000 | 6.2 | 88.6 | 5.2 | 1.000 | 0.0 | 95.5 | 4.5 | 1.000 | 2.3 | 94.4 | 3.3 | 1.000 |
| MCHC (g/dL) | 25.3 | 72.7 | 2.1 | **<0.001** | 14.1 | 85.9 | 0.0 | **<0.001** | 18.2 | 81.8 | 0.0 | 0.930 | 14.0 | 85.6 | 0.5 | **<0.001** |
| RDW (%) | 5.7 | 90.2 | 4.1 | 1.000 | 5.2 | 90.3 | 4.5 | 1.000 | 13.6 | 86.4 | 0.0 | 1.000 | 16.7 | 80.9 | 2.3 | **<0.001** |
| PLT (10^3^/µL) | 5.7 | 86.6 | 7.7 | 1.000 | 2.1 | 90.0 | 7.9 | 0.548 | 4.5 | 90.9 | 4.5 | 1.000 | 2.3 | 86.0 | 11.6 | **0.001** |
| MPV (fL) | 1.5 | 95.9 | 2.6 | 1.000 | 1.7 | 94.8 | 3.4 | 1.000 | 0.0 | 90.9 | 9.1 | 1.000 | 1.4 | 94.4 | 4.2 | 1.000 |
| NE (%) | 18.6 | 78.4 | 3.1 | **<0.001** | 15.5 | 78.3 | 6.2 | **<0.001** | 9.1 | 90.9 | 0.0 | 1.000 | 18.6 | 75.3 | 6.0 | **<0.001** |
| LY (%) | 4.6 | 77.3 | 18.0 | **<0.001** | 7.2 | 82.1 | 10.7 | **<0.001** | 0.0 | 95.5 | 4.5 | 1.000 | 8.4 | 76.7 | 14.9 | **<0.001** |
| MO (%) | 1.0 | 92.3 | 6.7 | 1.000 | 3.1 | 89.3 | 7.6 | 1.000 | 0.0 | 90.9 | 9.1 | 1.000 | 0.5 | 93.5 | 6.0 | 0.657 |
| EO (%) | 8.8 | 90.7 | 0.5 | 0.103 | 8.6 | 90.7 | 0.7 | **0.008** | 4.5 | 86.4 | 9.1 | 1.000 | 11.6 | 87.0 | 1.4 | **<0.001** |
| BA (%) | 1.0 | 94.8 | 4.1 | 1.000 | 0.7 | 93.8 | 5.5 | 0.268 | 0.0 | 100.0 | 0.0 | 1.000 | 0.0 | 94.4 | 5.6 | 0.273 |
| NE# (10^3^/µL) | 11.3 | 82.5 | 6.2 | 0.014 | 7.2 | 84.1 | 8.6 | 0.239 | 4.5 | 90.9 | 4.5 | 1.000 | 12.6 | 80.5 | 7.0 | **<0.001** |
| LY# (10^3^/µL) | 3.1 | 83.0 | 13.9 | **<0.001** | 2.8 | 88.3 | 9.0 | 0.186 | 0.0 | 100.0 | 0.0 | 1.000 | 0.9 | 89.3 | 9.8 | **0.017** |
| MO# (10^3^/µL) | 0.0 | 82.5 | 17.5 | **<0.001** | 2.1 | 77.9 | 20.0 | **<0.001** | 0.0 | 95.5 | 4.5 | 1.000 | 1.4 | 84.2 | 14.4 | **<0.001** |
| EO# (10^3^/µL) | 13.9 | 84.5 | 1.5 | **<0.001** | 12.1 | 86.9 | 1.0 | **<0.001** | 18.2 | 77.3 | 4.5 | 1.000 | 12.6 | 86.5 | 0.9 | **<0.001** |
| BA# (10^3^/µL) | 0.5 | 91.2 | 8.2 | 0.206 | 0.3 | 91.4 | 8.3 | **0.006** | 0.0 | 100.0 | 0.0 | 1.000 | 0.0 | 93.0 | 7.0 | 0.143 |

**Table S2. Multivariable model of disease-relevant features with residual age in T1D individuals.** Standardized regression estimates and standard errors are shown for continuous covariates, and the Type II ANOVA p-values are shown for categorical covariates with p-values <0.05 shown in bold. (See also Figure 3C)

| **Disease-Relevant Features** | **Estimate** | **Std. Error** | **p-value** |
| --- | --- | --- | --- |
| **Continuous features** |  |  |  |
| T1D Diagnosis Age | 0.878 | 0.559 | 0.120 |
| T1D Duration | 1.470 | 0.483 | **0.003** |
| BMI %tile | 1.752 | 0.468 | **<0.001** |
| HbA1C | 0.495 | 0.475 | 0.301 |
| Rested Blood Glucose | 1.102 | 0.494 | **0.029** |
| GRS1 | -0.236 | 0.540 | 0.663 |
| **Categorical features** |  |  |  |
| Sex | – | – | 0.228 |
| Ethnicity | – | – | 0.606 |
| Race | – | – | 0.957 |

**Table S3. Cellular subset frequencies and phenotypes assessed in peripheral blood of T1D and controls.** Outcome measures from flow cytometry (gated cell population frequency, mean fluorescence intensity (mfi), or normalized marker intensity (index) of (indicated parent population)) or complete blood count (CBC) assessment were compared in T1D and control subjects. Testing was performed using a non-parametric Kruskal Wallis followed by a post-hoc Dunn’s test with a Benjamini-Hochberg multiplicity adjustment. (See also Figure 4C)

| **Cell frequency or phenotype (Parent population)** | **Definition** | **Difference** | **Mean Quantile Difference** | **p-value** |
| --- | --- | --- | --- | --- |
| CXCR3^lo^ (CD8 Naïve) | CD183^lo^ % of CD3^+^CD4^-^ Naive | T1D ↑ | 0.21 | <0.001 |
| CXCR3^lo^ (CD8 CD45RO^-^) | CD183^lo^ % of CD3^+^CD4^-^CD45RO^-^ | T1D ↑ | 0.18 | <0.001 |
| CXCR3^+^ (Naive CD8) | CD183^+^ % of Naive CD8 | T1D ↑ | 0.142 | <0.001 |
| Naive (CD8) | Naive % of CD8 | T1D ↑ | 0.113 | <0.001 |
| Hemoglobin | HGB (g/dL) | T1D ↑ | 0.112 | 0.001 |
| CD123^+^ (MNCs) | CD123^+^ % of MNCs [mononuclear cells] | T1D ↑ | 0.11 | <0.001 |
| Hematocrit | HCT (%) | T1D ↑ | 0.108 | 0.003 |
| PD1^+^ (Naive CD4 ) | CD279^+^ % of Naive CD4 | T1D ↑ | 0.107 | <0.001 |
| CXCR3^lo^ (CD8 Temra) | CD183^lo^ % of CD3^+^CD4^-^ Temra | T1D ↑ | 0.101 | 0.001 |
| Activated Memory (Treg) | CD45RO^+^HLADR^+^CCR4^+^ % of Treg | T1D ↑ | 0.098 | 0.002 |
| Memory Th2 (Tconv) | CD45RO^+^CCR4^+^ % of Tconv | T1D ↑ | 0.096 | 0.003 |
| MCV | MCV [mean corpuscular volume] (fL) | T1D ↑ | 0.092 | 0.009 |
| Granulocyte (Leukocytes) | Granulocyte % of Leukocytes | T1D ↑ | 0.09 | 0.002 |
| MCH | MCH (pg) | T1D ↑ | 0.089 | 0.011 |
| PD1^+^ (CD8 Temra) | CD279^+^ % of CD8 Temra | T1D ↑ | 0.089 | 0.009 |
| HLA-DR mfi (Monocytes) | HLA-DR mfi of Monocytes | T1D ↑ | 0.088 | 0.014 |
| CXCR3^+^ (Tfh) | CD183^+^ % of Tfh | T1D ↑ | 0.087 | 0.002 |
| PD1^+^ (Naive CD8) | CD279^+^ % of Naive CD8 | T1D ↑ | 0.085 | 0.009 |
| CD38^+^HLADR^-^ (CD8) | CD38^+^HLADR^-^ % of CD8 | T1D ↑ | 0.084 | 0.01 |
| HLADR^+^CD38^+^ (Memory Th17) | HLADR^+^CD38^+^ % of CD4^+^CD45RO^+^CD183^-^CD196^+^ | T1D ↑ | 0.078 | 0.024 |
| Absolute platelet count | PLT (10^3^/µL) | T1D ↑ | 0.076 | 0.019 |
| Tcm (CD4) | Tcm % of CD4 | T1D ↑ | 0.074 | 0.027 |
| CXCR5^+^ (Naive CD4) | CD185^+^ % of Naive CD4 | T1D ↑ | 0.073 | 0.028 |
| Memory (Treg) | CD45RO^+^ % of Treg | T1D ↑ | 0.071 | 0.034 |
| CD45RO^+^CCR4^+^ (CD8) | CD45RO^+^CCR4^+^ % of CD3^+^CD4^-^ | T1D ↑ | 0.069 | 0.016 |
| CD56^bright^ (NK) | CD56^bright^ % of NK | T1D ↑ | 0.067 | 0.03 |
| PD1 index (Tcm CD4) | Tcm CD4 CD279mfi Rel. Diff. | T1D ↓ | 0.054 | 0.041 |
| NK (Lymphocytes) | NK % of Lymphocytes | T1D ↓ | 0.064 | 0.048 |
| CD38^-^HLADR^-^ (Naive CD8) | CD38^-^HLADR^-^ % of Naive CD8 | T1D ↓ | 0.066 | 0.023 |
| CD56dim (NK) | CD56dim % of NK | T1D ↓ | 0.069 | 0.028 |
| PD1 index (Tem CD4) | Tem CD4 CD279mfi Rel. Diff. | T1D ↓ | 0.07 | 0.035 |
| Lymphocytes (CBC) | LY (%) | T1D ↓ | 0.071 | 0.036 |
| MNC (Leukocytes) | MNC % of Leukocytes | T1D ↓ | 0.073 | 0.021 |
| CD38^+^HLADR^-^ (CD8 Temra) | CD38^+^HLADR^-^ % of CD8 Temra | T1D ↓ | 0.075 | 0.023 |
| CD38^-^HLADR^-^ (Naive CD4) | CD38^-^HLADR^-^ % of Naive CD4 | T1D ↓ | 0.076 | 0.022 |
| CXCR3^hi^ (CD8 CD45RO^-^) | CD183^hi^ % of CD3^+^CD4^-^CD45RO^-^ | T1D ↓ | 0.08 | 0.007 |
| Transitional (B cells) | Transitional % of B cells | T1D ↓ | 0.09 | 0.005 |
| PD1mfi (Temra CD4) | Temra CD4 CD279mfi | T1D ↓ | 0.091 | 0.004 |
| PD1mfi (Tcm CD4) | Tcm CD4 CD279mfi | T1D ↓ | 0.092 | 0.001 |
| CD38^-^HLADR^-^ (CD8) | CD38^-^HLADR^-^ % of CD8 | T1D ↓ | 0.093 | 0.004 |
| DN (T cells) | DN % of T cells | T1D ↓ | 0.106 | 0.001 |
| PD1mfi (Tcm CD8) | Tcm CD8 CD279mfi | T1D ↓ | 0.108 | <0.001 |
| CXCR3^-^ (CD8 Temra) | CD183^-^ % of CD3^+^CD4^-^ Temra | T1D ↓ | 0.108 | <0.001 |
| Tem (CD8) | Tem % of CD8 | T1D ↓ | 0.112 | <0.001 |
| PD1mfi (Tem CD4) | Tem CD4 CD279mfi | T1D ↓ | 0.117 | <0.001 |
| RDW | RDW (%) | T1D ↓ | 0.154 | <0.001 |
| CXCR3^-^ (CD8 CD45RO^-^ ) | CD183^-^ % of CD3^+^CD4^-^CD45RO^-^ | T1D ↓ | 0.163 | <0.001 |
| CXCR3^-^ (CD8 Naïve) | CD183^-^ %of CD3^+^CD4^-^ Naive | T1D ↓ | 0.189 | <0.001 |
| Naive (B cells) | Naive % of B cells | T1D ↑ | 0.066 | ns |
| Myeloid (DCs) | Myeloid % of DCs | T1D ↑ | 0.065 | ns |
| HLA-DR mfi (DCs) | HLA-DR mfi of DCs | T1D ↑ | 0.065 | ns |
| CXCR5^+^ (CD8 Temra) | CD185^+^ % of CD8 Temra | T1D ↑ | 0.064 | ns |
| Neutrophils (CBC) | NE (%) | T1D ↑ | 0.062 | ns |
| CXCR3^+^ (CD4 Tcm) | CD183^+^ % of CD4 Tcm | T1D ↑ | 0.06 | ns |
| CD38^+^HLADR^-^ (Naive CD4) | CD38^+^HLADR^-^ % of Naive CD4 | T1D ↑ | 0.06 | ns |
| CD38^+^HLADR^-^ (Naive CD8) | CD38^+^HLADR^-^ % of Naive CD8 | T1D ↑ | 0.058 | ns |
| CXCR3^+^ (CD8 Temra) | CD183^+^ % of CD8 Temra | T1D ↑ | 0.055 | ns |
| CD38^-^HLADR^+^ (CD8 Temra) | CD38^-^HLADR^+^ % of CD8 Temra | T1D ↑ | 0.054 | ns |
| RBC count | RBC (10^6^/µL) | T1D ↑ | 0.054 | ns |
| Activated Memory (CD8) | CD45RO^+^HLADR^+^CCR4^+^ % of CD3^+^CD4^-^ | T1D ↑ | 0.054 | ns |
| CXCR3^lo^ (CD8 Tcm) | CD183^lo^ % of CD3^+^CD4^-^ Tcm | T1D ↑ | 0.053 | ns |
| MCHC | MCHC (g/dL) | T1D ↑ | 0.053 | ns |
| CD38^-^HLADR^-^ (CD8 Temra) | CD38^-^HLADR^-^ % of CD8 Temra | T1D ↑ | 0.052 | ns |
| CXCR5^+^ (CD8 Tem) | CD185^+^ % of CD8 Tem | T1D ↑ | 0.051 | ns |
| PD1^+^ (CD8 Tem) | CD279^+^ % of CD8 Tem | T1D ↑ | 0.051 | ns |
| CXCR3^+^ (Naive CD4) | CD183^+^ % of Naive CD4 | T1D ↑ | 0.051 | ns |
| HLADR^+^ (CD8CD45RO^+^CXCR3^-^CCR6^-^) | HLA^+^ % of CD4^-^CD45RO^+^CD183^-^CD196^-^ | T1D ↑ | 0.05 | ns |
| CXCR5^+^ (CD4 Temra) | CD185^+^ % of CD4 Temra | T1D ↑ | 0.049 | ns |
| Memory Th2 (CD4) | CD45RO^+^CD183^-^CD196^-^ % of CD3^+^CD4^+^ | T1D ↑ | 0.048 | ns |
| PD1 index (Temra CD8) | Temra CD8 CD279mfi Rel. Diff. | T1D ↑ | 0.048 | ns |
| Basophils (CBC) | BA (%) | T1D ↑ | 0.047 | ns |
| CD4 (T cells) | CD4 % of T cells | T1D ↑ | 0.043 | ns |
| HLADR^+^ (Memory Th17) | HLADR^+^ % of CD4^+^CD45RO^+^CD183^-^CD196^+^ | T1D ↑ | 0.043 | ns |
| Memory Th1/17 (CD4) | CD45RO^+^CD183^+^CD196^+^ % of CD3^+^CD4^+^ | T1D ↑ | 0.042 | ns |
| Activated Memory (Tconv) | CD45RO^+^HLADR^+^CCR4^+^ % of Tconv | T1D ↑ | 0.041 | ns |
| CXCR3^hi^ (CD8 Temra) | CD183^hi^ % of CD3^+^CD4^-^ Temra | T1D ↑ | 0.041 | ns |
| Tfh (CD4^+^CD45RA^-^CXCR5^+^) | Tfh % of CD4^+^CD45RA^-^CD185^+^ | T1D ↑ | 0.041 | ns |
| HLADR^+^CD38^+^ (Memory Th1/17) | HLADR^+^CD38^+^ % of CD4^+^CD45RO^+^CD183^+^CD196^+^ | T1D ↑ | 0.036 | ns |
| CXCR3^lo^ (CD8 Tem) | CD183^lo^ % of CD3^+^CD4^-^ Tem | T1D ↑ | 0.034 | ns |
| T cells (Lymphocytes) | T cells % of Lymphocytes | T1D ↑ | 0.034 | ns |
| Non-Class-switched Memory (B cells) | Non-Class-switched Memory % of B cells | T1D ↑ | 0.034 | ns |
| Tcm (CD8) | Tcm % of CD8 | T1D ↑ | 0.032 | ns |
| Memory Th17 (CD4) | CD45RO^+^CD183^-^CD196^+^ % of CD3^+^CD4^+^ | T1D ↑ | 0.031 | ns |
| Absolute neutrophil count | NE# (10^3^/µL) | T1D ↑ | 0.031 | ns |
| Memory Th1 (CD4) | CD45RO^+^CD183^+^CD196^-^ % of CD3^+^CD4^+^ | T1D ↑ | 0.03 | ns |
| Classical (Monocytes) | Classical % of Monocytes | T1D ↑ | 0.029 | ns |
| HLADR^+^ (Memory Th1/17) | HLADR^+^ % of CD4^+^CD45RO^+^CD183^+^CD196^+^ | T1D ↑ | 0.027 | ns |
| CD38^-^HLADR^+^ (CD4 Temra) | CD38^-^HLADR^+^ % of CD4 Temra | T1D ↑ | 0.027 | ns |
| HLADR^+^CD38^+^ (CD8CD45RO^+^CXCR3^-^CCR6^-^) | HLA^+^CD38^+^ % of CD4^-^CD45RO^+^CD183^-^CD196^-^ | T1D ↑ | 0.026 | ns |
| Monocytes (MNCs) | Monocytes % of MNCs | T1D ↑ | 0.024 | ns |
| CD38^+^HLADR^+^ (Naive CD8) | CD38^+^HLADR^+^ % of Naive CD8 | T1D ↑ | 0.024 | ns |
| CD38^-^HLADR^-^ (CD4 Tem) | CD38^-^HLADR^-^ % of CD4 Tem | T1D ↑ | 0.023 | ns |
| Tfh (CD4) | Tfh % of CD4 | T1D ↑ | 0.023 | ns |
| CD8 (T cells) | CD8 % of T cells | T1D ↑ | 0.022 | ns |
| CXCR5^+^ (CD4 Tem) | CD185^+^ % of CD4 Tem | T1D ↑ | 0.02 | ns |
| PD1 index (Tem CD8) | Tem CD8 CD279mfi Rel. Diff. | T1D ↑ | 0.019 | ns |
| Absolute basophil count | BA# (10^3^/µL) | T1D ↑ | 0.017 | ns |
| CD38^+^HLADR^+^ (CD8) | CD38^+^HLADR^+^ % of CD8 | T1D ↑ | 0.017 | ns |
| Precursor Tfh (CD4^+^CD45RA^-^CXCR5^+^) | Precursor Tfh % of CD4^+^CD45RA^-^CD185^+^ | T1D ↑ | 0.017 | ns |
| Tem (CD4) | Tem % of CD4 | T1D ↑ | 0.017 | ns |
| CD38^-^HLADR^-^ (CD4) | CD38^-^HLADR^-^ % of CD4 | T1D ↑ | 0.017 | ns |
| CD25mfi (CD45RO^-^ Tconv) | CD25mfi of CD45RO^-^ Tconv | T1D ↑ | 0.017 | ns |
| CD38^-^HLADR^+^ (CD8 Tcm) | CD38^-^HLADR^+^ % of CD8 Tcm | T1D ↑ | 0.016 | ns |
| HLADR^+^ (CD8CD45RO^+^CXCR3^+^CCR6^-^) | HLA^+^ % of CD4^-^CD45RO^+^CD183^+^CD196^-^ | T1D ↑ | 0.016 | ns |
| CD38^+^HLADR^-^ (CD4 Tcm) | CD38^+^HLADR^-^ % of CD4 Tcm | T1D ↑ | 0.015 | ns |
| PD1^+^ (CD4 Tcm) | CD279^+^ % of CD4 Tcm | T1D ↑ | 0.015 | ns |
| DC (MNCs) | DC % of MNCs | T1D ↑ | 0.015 | ns |
| CD38^+^HLADR^+^ (CD4 Temra) | CD38^+^HLADR^+^ % of CD4 Temra | T1D ↑ | 0.014 | ns |
| CD38^+^HLADR^+^ (CD8 Tem) | CD38^+^HLADR^+^ % of CD8 Tem | T1D ↑ | 0.014 | ns |
| HLADR^+^CD38^+^ (Memory Th2) | HLADR^+^CD38^+^ % of CD4^+^CD45RO^+^CD183^-^CD196^-^ | T1D ↑ | 0.013 | ns |
| CD38^-^HLADR^-^ (CD4 Temra) | CD38^-^HLADR^-^ % of CD4 Temra | T1D ↑ | 0.012 | ns |
| HLADR^+^CD38^+^ (CD8CD45RO^+^CXCR3^+^CCR6^-^) | HLA^+^CD38^+^ % of CD4^-^CD45RO^+^CD183^+^CD196^-^ | T1D ↑ | 0.011 | ns |
| Precursor Tfh (CD4) | Precursor Tfh % of CD4 | T1D ↑ | 0.009 | ns |
| CD38^-^HLADR^+^ (CD8 Tem) | CD38^-^HLADR^+^ % of CD8 Tem | T1D ↑ | 0.009 | ns |
| CXCR3^+^ (CD4 Tem) | CD183^+^ % of CD4 Tem | T1D ↑ | 0.008 | ns |
| CD38^+^HLADR^+^ (CD4 Tem) | CD38^+^HLADR^+^ % of CD4 Tem | T1D ↑ | 0.007 | ns |
| CD38^+^HLADR^+^ (Naive CD4) | CD38^+^HLADR^+^ % of Naive CD4 | T1D ↑ | 0.006 | ns |
| B cells (Lymphocytes) | B cells % of Lymphocytes | T1D ↑ | 0.005 | ns |
| WBC count | WBC (10^3^/µL) | T1D ↑ | 0.005 | ns |
| CXCR3^+^ (CD4 Temra) | CD183^+^ % of CD4 Temra | T1D ↑ | 0.005 | ns |
| CD38^-^HLADR^+^ (CD4 Tcm) | CD38^-^HLADR^+^ % of CD4 Tcm | T1D ↑ | 0.005 | ns |
| CD25mfi (Naïve CD8) | CD25mfi of CD45RO^-^CD3^+^CD4^-^ | T1D ↑ | 0.005 | ns |
| CXCR3^+^ (CD8 Tcm) | CD183^+^ % of CD8 Tcm | T1D ↑ | 0.004 | ns |
| CD38^-^HLADR^+^ (Naive CD8) | CD38^-^HLADR^+^ % of Naive CD8 | T1D ↑ | 0.004 | ns |
| HLADR^+^CD38^+^ (CD8CD45RO^+^CXCR3^lo^CCR6^+^) | HLA^+^CD38^+^ % of CD4^-^CD45RO^+^CD183^lo^CD196^+^ | T1D ↑ | 0.003 | ns |
| CD38^-^HLADR^+^ (CD4) | CD38^-^HLADR^+^ % of CD4 | T1D ↑ | 0.003 | ns |
| CD25mfi (Memory CD8) | CD25mfi of CD45RO^+^CD3^+^CD4^-^ | T1D ↑ | 0.003 | ns |
| CD38^+^HLADR^+^ (CD4) | CD38^+^HLADR^+^ % of CD4 | T1D ↑ | 0.001 | ns |
| CD38^-^HLADR^+^ (CD4 Tem) | CD38^-^HLADR^+^ % of CD4 Tem | T1D ↑ | 0.001 | ns |
| PD1^+^ (CD4 Temra) | CD279^+^ % of CD4 Temra | T1D ↑ | 0 | ns |
| CXCR3^lo^CCR6^+^ (CD8) | CD183^lo^CD196^+^ % of CD3^+^CD4^-^ | T1D ↑ | 0 | ns |
| CD38^+^HLADR^+^ (CD8 Tcm) | CD38^+^HLADR^+^ % of CD8 Tcm | T1D ↑ | 0 | ns |
| CD38^-^HLADR^-^ (CD8 Tem) | CD38^-^HLADR^-^ % of CD8 Tem | T1D ↓ | 0.001 | ns |
| HLADR^+^CD38^+^ (Memory Th1) | HLADR^+^CD38^+^ % of CD4^+^CD45RO^+^CD183^+^CD196^-^ | T1D ↓ | 0.001 | ns |
| DP (T cells) | DP % of T cells | T1D ↓ | 0.002 | ns |
| CD38^+^HLADR^-^ (CD8 Tcm) | CD38^+^HLADR^-^ % of CD8 Tcm | T1D ↓ | 0.002 | ns |
| Treg (CD4) | Treg % of CD4^+^ | T1D ↓ | 0.003 | ns |
| Conventional (DCs) | CD11c^-^CD123^-^ % of DC | T1D ↓ | 0.006 | ns |
| CXCR5^+^ (Naive CD8) | CD185^+^ % of Naive CD8 | T1D ↓ | 0.006 | ns |
| CD16^+^ (CD56^bright^ NK) | CD16^+^ % of CD56^bright^ NK | T1D ↓ | 0.007 | ns |
| PD1mfi (Temra CD8) | Temra CD8 CD279mfi | T1D ↓ | 0.008 | ns |
| Absolute monocyte count | MO# (10^3^/µL) | T1D ↓ | 0.009 | ns |
| CD38^+^HLADR^+^ (CD8 Temra) | CD38^+^HLADR^+^ % of CD8 Temra | T1D ↓ | 0.011 | ns |
| Memory Treg CD25 index | CD45RO^+^ Treg CD25mfi Rel. Diff. | T1D ↓ | 0.013 | ns |
| CD38^+^HLADR^+^ (CD4 Tcm) | CD38^+^HLADR^+^ % of CD4 Tcm | T1D ↓ | 0.014 | ns |
| HLADR^+^ (CD8CD45RO^+^CXCR3^lo^CCR6^+^) | HLA^+^ % of CD4^-^CD45RO^+^CD183^lo^CD196^+^ | T1D ↓ | 0.014 | ns |
| CXCR5^+^ (CD4 Tcm) | CD185^+^ % of CD4 Tcm | T1D ↓ | 0.015 | ns |
| CXCR3^-^ (CD8 Tem) | CD183^-^ % of CD3^+^CD4^-^ Tem | T1D ↓ | 0.015 | ns |
| CXCR5^+^ (CD8 Tcm) | CD185^+^ % of CD8 Tcm | T1D ↓ | 0.016 | ns |
| CD25mfi (CD45RO^+^ Tconv | CD25mfi of CD45RO^+^ Tconv | T1D ↓ | 0.016 | ns |
| CD38^+^HLADR^-^ (CD4) | CD38^+^HLADR^-^ % of CD4 | T1D ↓ | 0.017 | ns |
| CXCR3^+^ (CD8) | CD183^+^CD196^-^ % of CD3^+^CD4^-^ | T1D ↓ | 0.018 | ns |
| PD1^+^ (CD8 Tcm) | CD279^+^ % of CD8 Tcm | T1D ↓ | 0.018 | ns |
| Memory CD8 CD25 index | CD45RO^+^CD3^+^CD4^-^ CD25mfi Rel. Diff. | T1D ↓ | 0.019 | ns |
| CD38^-^HLADR^-^ (CD8 Tcm) | CD38^-^HLADR^-^ % of CD8 Tcm | T1D ↓ | 0.02 | ns |
| HLADR^+^ (Memory Th2) | HLADR^+^ % of CD4^+^CD45RO^+^CD183^-^CD196^-^ | T1D ↓ | 0.022 | ns |
| HLADR^+^ (Memory Th1) | HLADR^+^ % of CD4^+^CD45RO^+^CD183^+^CD196^-^ | T1D ↓ | 0.022 | ns |
| CD38^-^HLADR^-^ (CD4 Tcm) | CD38^-^HLADR^-^ % of CD4 Tcm | T1D ↓ | 0.024 | ns |
| Memory Tconv CD25 index | CD45RO^+^ Tconv CD25mfi Rel. Diff. | T1D ↓ | 0.024 | ns |
| CD38^-^HLADR^+^ (CD8) | CD38^-^HLADR^+^ % of CD8 | T1D ↓ | 0.026 | ns |
| CXCR3^-^ (CD8 Tcm) | CD183^-^ % of CD3^+^CD4^-^ Tcm | T1D ↓ | 0.027 | ns |
| CXCR3^hi^ (CD8 Tem) | CD183^hi^ % of CD3^+^CD4^-^ Tem | T1D ↓ | 0.027 | ns |
| Memory Tfh (CD4) | Memory Tfh % of CD4 | T1D ↓ | 0.028 | ns |
| Non-Classical (Monocytes) | Non-Classical % of Monocytes | T1D ↓ | 0.029 | ns |
| CXCR3^hi^ (CD8 Tcm) | CD183^hi^ % of CD3^+^CD4^-^ Tcm | T1D ↓ | 0.029 | ns |
| Absolute eosinophil count | EO# (10^3^/µL) | T1D ↓ | 0.032 | ns |
| Memory Tfh (CD4^+^CD45RA^-^CXCR5^+^) | Memory Tfh % of CD4^+^CD45RA^-^CD185^+^ | T1D ↓ | 0.032 | ns |
| Eosinophils (CBC) | EO (%) | T1D ↓ | 0.032 | ns |
| PD1mfi (Tem CD8) | Tem CD8 CD279mfi | T1D ↓ | 0.032 | ns |
| PD1^+^ (CD4 Tem) | CD279^+^ % of CD4 Tem | T1D ↓ | 0.034 | ns |
| Temra (CD4) | Temra % of CD4 | T1D ↓ | 0.036 | ns |
| Monocytes (CBC) | MO (%) | T1D ↓ | 0.036 | ns |
| PD1 index (Temra CD4) | Temra CD4 CD279mfi Rel. Diff. | T1D ↓ | 0.039 | ns |
| MPV | MPV [mean platelet volume] (fL) | T1D ↓ | 0.039 | ns |
| Class-switched Memory (B cells) | Class-switched Memory % of B cells | T1D ↓ | 0.04 | ns |
| CXCR3^+^ (CD8 Tem) | CD183^+^ % of CD8 Tem | T1D ↓ | 0.041 | ns |
| CXCR3^hi^ (CD8 Naïve) | CD183^hi^ % of CD3^+^CD4^-^ Naive | T1D ↓ | 0.041 | ns |
| CD38^+^HLADR^-^ (CD8 Tem) | CD38^+^HLADR^-^ % of CD8 Tem | T1D ↓ | 0.042 | ns |
| CD38^-^HLADR^+^ (Naive CD4) | CD38^-^HLADR^+^ % of Naive CD4 | T1D ↓ | 0.043 | ns |
| Naive (CD4) | Naive % of CD4 | T1D ↓ | 0.047 | ns |
| CXCR3^-^CCR6^-^ (CD8) | CD183^-^CD196^-^ % of CD3^+^CD4^-^ | T1D ↓ | 0.048 | ns |
| CD38^+^HLADR^-^ (CD4 Tem) | CD38^+^HLADR^-^ % of CD4 Tem | T1D ↓ | 0.049 | ns |
| PD1 index (Tcm CD8) | Tcm CD8 CD279mfi Rel. Diff. | T1D ↓ | 0.05 | ns |
| CD38^+^HLADR^-^ (CD4 Temra) | CD38^+^HLADR^-^ % of CD4 Temra | T1D ↓ | 0.052 | ns |
| PD1mfi (Naive CD4) | Naive CD4 CD279mfi | T1D ↓ | 0.053 | ns |
| Temra (CD8) | Temra % of CD8 | T1D ↓ | 0.055 | ns |
| Plasmacytoid (DCs) | Plasmacytoid % DCs | T1D ↓ | 0.055 | ns |
| PD1mfi (Naive CD8) | Naive CD8 CD279mfi | T1D ↓ | 0.056 | ns |
| CD25mfi (Memory Treg) | CD25mfi of CD45RO^+^ Treg | T1D ↓ | 0.057 | ns |
| CD25mfi (Naïve Treg) | CD25mfi of CD45RO^-^ Treg | T1D ↓ | 0.059 | ns |
| Absolute lymphocyte count | LY# (10^3^/µL) | T1D ↓ | 0.06 | ns |
| Plasmablast (B cells) | Plasmablast/Plasma Cell % of B cells | T1D ↓ | 0.065 | ns |

**Table S4. Logistic regression of Genotype-Tissue Expression project (GTeX) *PDCD1* expression quantitative trait loci (eQTL) in CTR versus T1D subjects.** Association between single nucleotide polymorphisms (SNP) and clinical status tested with sex and population stratification as covariates. A1 = effect allele, A2 = non-effect allele, OR = odds ratio, STAT = coefficient t-statistic, P = p-value. (See also Figure S14)

| **SNP** | **A1** | **A2** | **OR** | **STAT** | **P** |
| --- | --- | --- | --- | --- | --- |
| rs6422701 | C | T | 0.627 | -2.431 | 0.015 |
| rs35905226 | T | C | 1.616 | 2.344 | 0.019 |
| rs34623950 | G | A | 1.501 | 2.019 | 0.044 |
| rs28435574 | A | G | 1.454 | 1.911 | 0.056 |
| rs6734491 | T | C | 1.443 | 1.865 | 0.062 |
| rs35380510 | T | C | 1.401 | 1.637 | 0.102 |
| rs11683104 | C | T | 1.345 | 1.486 | 0.137 |
| rs67309595 | A | G | 1.331 | 1.458 | 0.145 |
| rs4596012 | C | T | 0.754 | -1.424 | 0.155 |
| rs35301767 | A | G | 0.750 | -1.322 | 0.186 |
| rs73108258 | A | G | 0.649 | -1.112 | 0.266 |
| rs66492541 | A | G | 1.169 | 0.787 | 0.432 |
| rs6422702 | A | G | 0.846 | -0.780 | 0.436 |
| rs78280775 | C | T | 0.890 | -0.573 | 0.567 |
| rs10755057 | T | G | 0.928 | -0.364 | 0.716 |
| rs35350837 | A | G | 0.937 | -0.322 | 0.748 |
| rs1106416 | T | C | 1.125 | 0.306 | 0.760 |
| rs79326616 | T | C | 0.917 | -0.300 | 0.764 |
| rs74898709 | A | G | 0.923 | -0.295 | 0.768 |
| rs78415814 | T | C | 0.918 | -0.287 | 0.774 |
| rs114496196 | C | A | 0.904 | -0.244 | 0.808 |
| rs74603454 | A | C | 0.948 | -0.196 | 0.845 |
| rs1106639 | A | G | 1.026 | 0.119 | 0.905 |
| rs10933569 | A | G | 1.003 | 0.014 | 0.989 |

**Table S5. QTL analysis.** Linear regression analysis of associations between 277 T1D genetic risk loci and 172 GAMLSS-corrected flow cytometric parameters, considering T1D status, sex, and population admixture as covariates. Beta coefficients are shown in the context of the T1D risk allele. False discovery rate (FDR) for total genotype and phenotype combinations determined by Benjamini-Hochberg method. Data are available in a supplementary Excel file. (See also Figure 7)

**Table S6. Reagents and resources used in the study.**

| **REAGENT or RESOURCE** | **SOURCE** | **IDENTIFIER** |
| --- | --- | --- |
| **Antibodies** | | |
| Pacific Blue anti-CD3 Pacific Blue anti-CD14 BV421 anti-CD197 FITC anti-CD4 AF488 anti-HLA-DR AF488 anti-IgD PerCP/Cy5.5 anti-CD4 PerCP/Cy5.5 anti-CD19 PerCP/Cy5.5 anti-CD45RA PerCP/Cy5.5 anti-CD123 PE anti-CD24 PE anti-CD56 PE anti-CD127 PE anti-CD183 PE anti-CD185 PE anti-CD197 PE/Cy7 anti-CD11c PE/Cy7 anti-CD27 PE/Cy7 anti-CD45RA PE/Cy7 anti-CD183 PE/Cy7 anti-CD194 PE/Cy7 anti-CD196 APC anti-CD16 APC anti-CD25 APC anti-CD38 APC anti-CD279 APC/H7 anti-CD3 APC/H7 anti-CD8 APC/H7 anti-CD19 APC/H7 anti-CD20 APC/H7 anti-CD45RO Human TruStain FcX™ True-Stain Monocyte Blocker | BioLegend BioLegend Biolegend Biolegend BioLegend BioLegend BioLegend BioLegend Biolegend Biolegend Biolegend Biolegend BioLegend BioLegend BioLegend BioLegend BioLegend BioLegend BioLegend BioLegend BioLegend BioLegend BioLegend BioLegend BioLegend eBioscience BD Biosciences BD Biosciences BD Biosciences BD Biosciences BD Biosciences BioLegend BioLegend | Cat#300417; RRID:AB_493094 Cat#301828; RRID:AB_2275670 Cat#353208; RRID:AB_10915137 Cat#300506; RRID:AB_314074 Cat#307620; RRID:AB_493175 Cat#348215; RRID:AB_11150397 Cat#300530; RRID:AB_893322 Cat#302230; RRID:AB_2073119 Cat#304122; RRID:AB_893357 Cat#306016; RRID:AB_2264693 Cat#311106; RRID:AB_314855 Cat#318306; RRID:AB_604101 Cat#351304; RRID:AB_10720185 Cat#353706; RRID:AB_10962912 Cat#356904; RRID:AB_2561813 Cat#353204; RRID:AB_10913813 Cat#337216; RRID:AB_2129790 Cat#302838; RRID:AB_2561919 Cat#304126; RRID:AB_10708879 Cat#353720; RRID:AB_11219383 Cat#359410; RRID:AB_2562431 Cat#353418; RRID:AB_10916518 Cat#302012; RRID:AB_314212 Cat#302610; RRID:AB_314280 Cat#356606; RRID:AB_2561902 Cat#17-2799-42; RRID:AB_11063701 Cat#560176; RRID:AB_1645475 Cat#560179; RRID:AB_1645481 Cat#560727; RRID:AB_1727437 Cat#560853; RRID:AB_10561681 Cat#561137; RRID:AB_10562194 Cat#422302; RRID:AB_2818986 Cat#426103 |
| **Critical Commercial Assays** | | |
| eBioscience 1-step Fix/Lyse Solution 10X Concentrate | ThermoFisher | Cat#00-5333-54 |
| **Software and Algorithms** | | |
| FlowJo v9 and v10 | BD Life Sciences | <https://www.flowjo.com> |

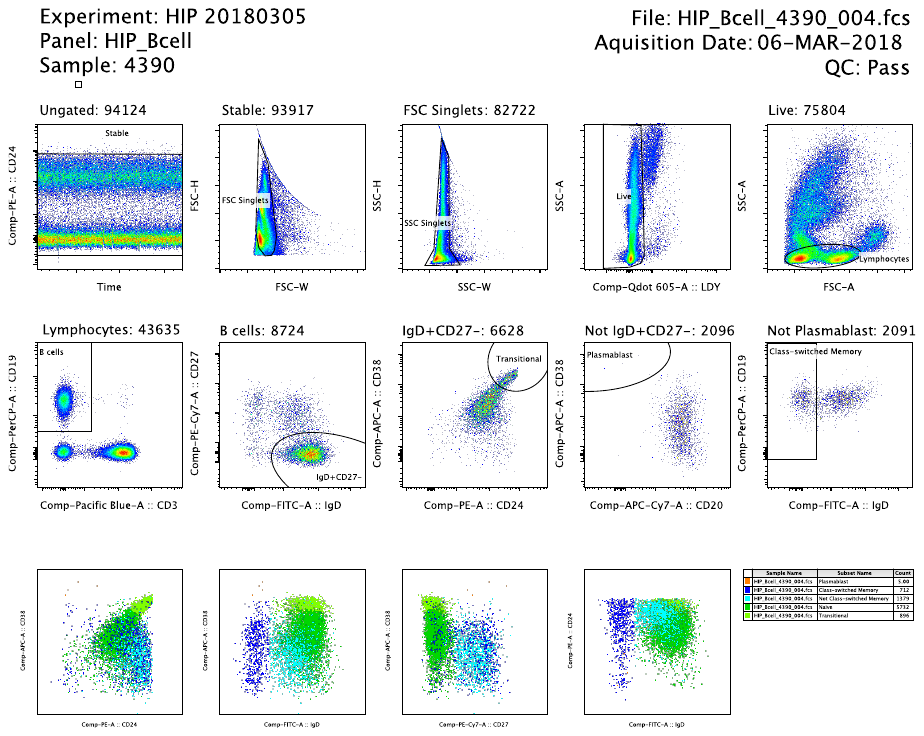
Figure S1. B cell panel gating. Gating based on time, forward scatter (FSC), and side scatter (SSC) was performed to select stably acquired singlets. The Qdot605 channel was used to remove autofluorescent granulocytes, then lymphocytes were gated based on FSC and SSC parameters. CD19^+^CD3^-^ B cells were subset into IgD^+^CD27^-^ cells, comprised of CD24^lo/-^CD38^lo/-^ naïve (dark green) and CD24^hi^CD38^hi^ transitional (light green) B cells. B cells that were not in the IgD^+^CD27^-^ gate were subset into CD20^-^CD38^hi^ plasmablasts (orange). Non-plasmablasts were designated as IgD^+^ non-class-switched (light blue) or IgD^-^ class-switched memory (dark blue) B cells.

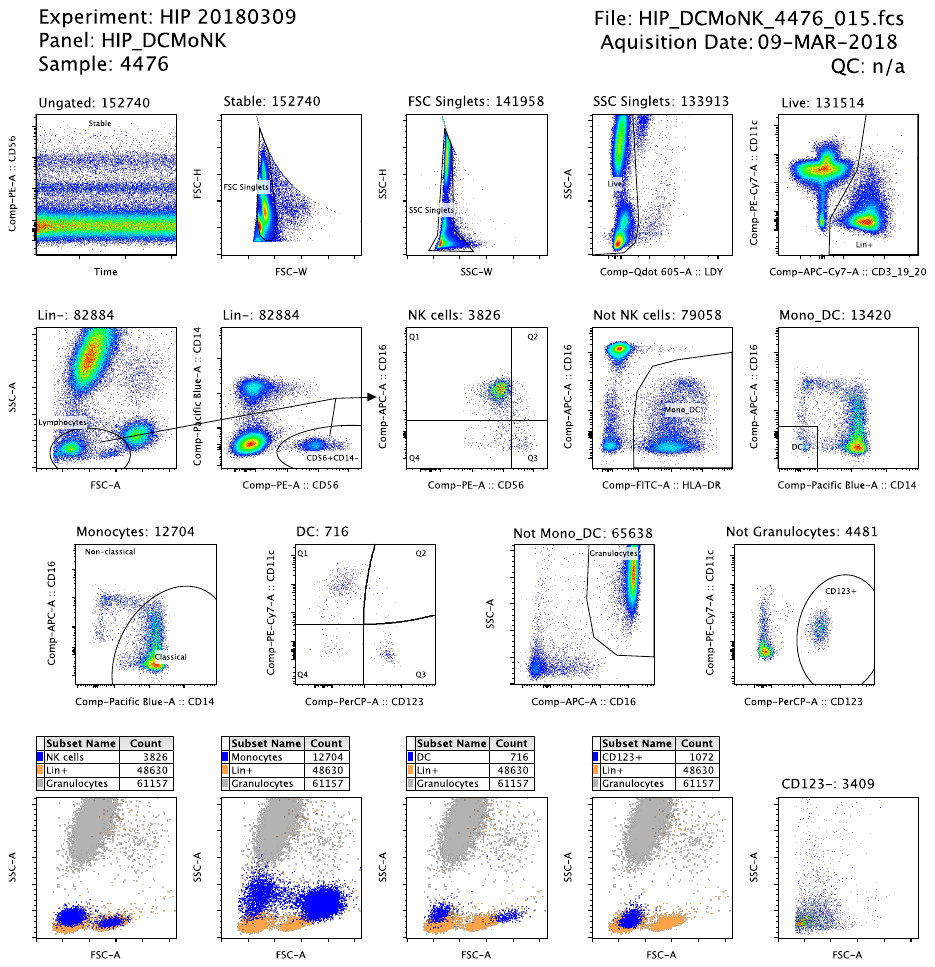
Figure S2. Innate immune cell panel gating. Gating based on time, forward scatter (FSC), and side scatter (SSC) was performed to select stably acquired singlets. The Qdot605/LDY channel was used to remove autofluorescent granulocytes, then lineage negative (Lin^-^) CD3^-^CD19^-^CD20^-^ cells were selected. Lymphocytes were gated based on FSC and SSC parameters, followed by gating on CD56^+^CD14^-^ natural killer (NK) cells (blue, bottom left). NK cells were quadrant gated into CD56^bright^, CD56^dim^, and CD16^+^ of CD56^bright^ subsets. HLA-DR^+^ non-NK cells were subset into CD14^-^CD16^-^ dendritic cells (DCs, blue, bottom middle), with non-DC as monocytes (blue, bottom second to left). Monocytes were grouped into CD14^+^CD16^-^ classical and CD14^dim^CD16^+^ non-classical subsets, while DC were grouped into CD11c^+^CD123^-^ myeloid, CD11c^-^CD123^+^ plasmacytoid, and CD11c^-^CD123^-^ conventional subsets. Non-monocyte/DC were gated for SSC^hi^CD16^hi^ granulocytes and non-granulocytes for other undefined CD123^+^ mononuclear cells (MNCs) (blue, bottom second to right). Size and granularity of previously defined major innate subsets compared to lineage positive (Lin^+^) cells (orange).

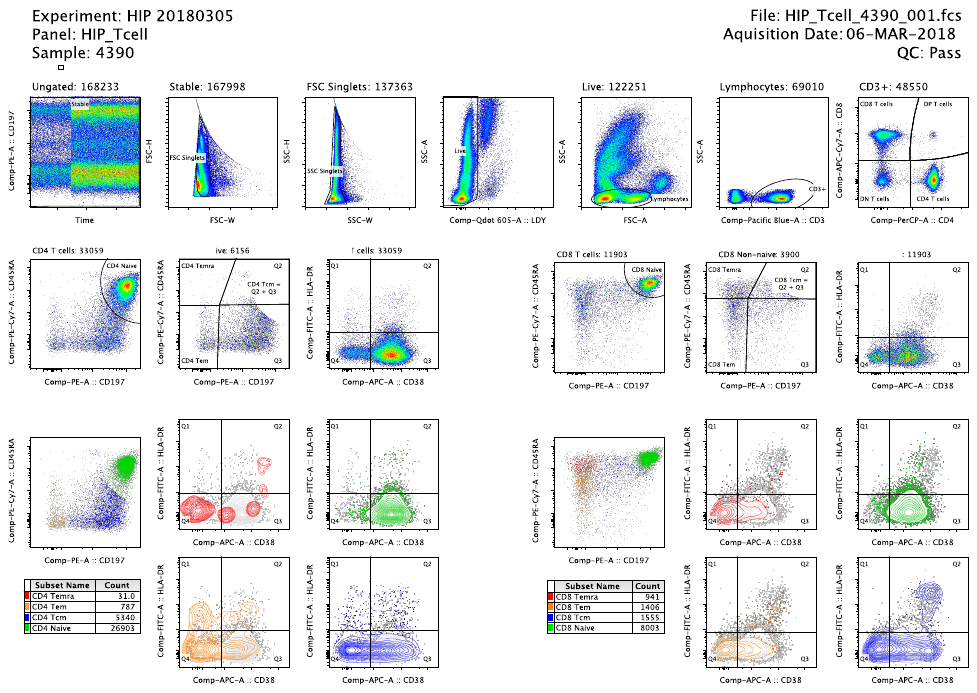
Figure S3. T cell panel gating. Gating based on time, forward scatter (FSC), and side scatter (SSC) was performed to select stably acquired singlets. The Qdot605/LDY channel was used to remove autofluorescent granulocytes, then lymphocytes were gated based on FSC and SSC parameters. CD3^+^ T cells were grouped into CD4^+^, CD8^+^, double negative (DN, CD4^-^CD8^-^), and double positive (DP, CD4^+^CD8^+^) T cell subsets. Gating was performed within CD4^+^ (left) and CD8^+^ (right) T cell populations to select CD45RA^+^CD197^+^ naïve (green), CD45RA^-^CD197^+^ central memory (Tcm, blue), CD45RA^-^CD197^-^ effector memory (Tem, orange), and CD45RA^+^CD197^-^ terminally differentiated effector memory (Temra, red) T cells. CD38^-^HLA-DR^-^, CD38^+^HLA-DR^-^, CD38^-^HLA-DR^+^, and CD38^+^HLA-DR^+^ subsets were gated within total as well as naïve, Tcm, Tem, and Temra CD4 or CD8 T cells.

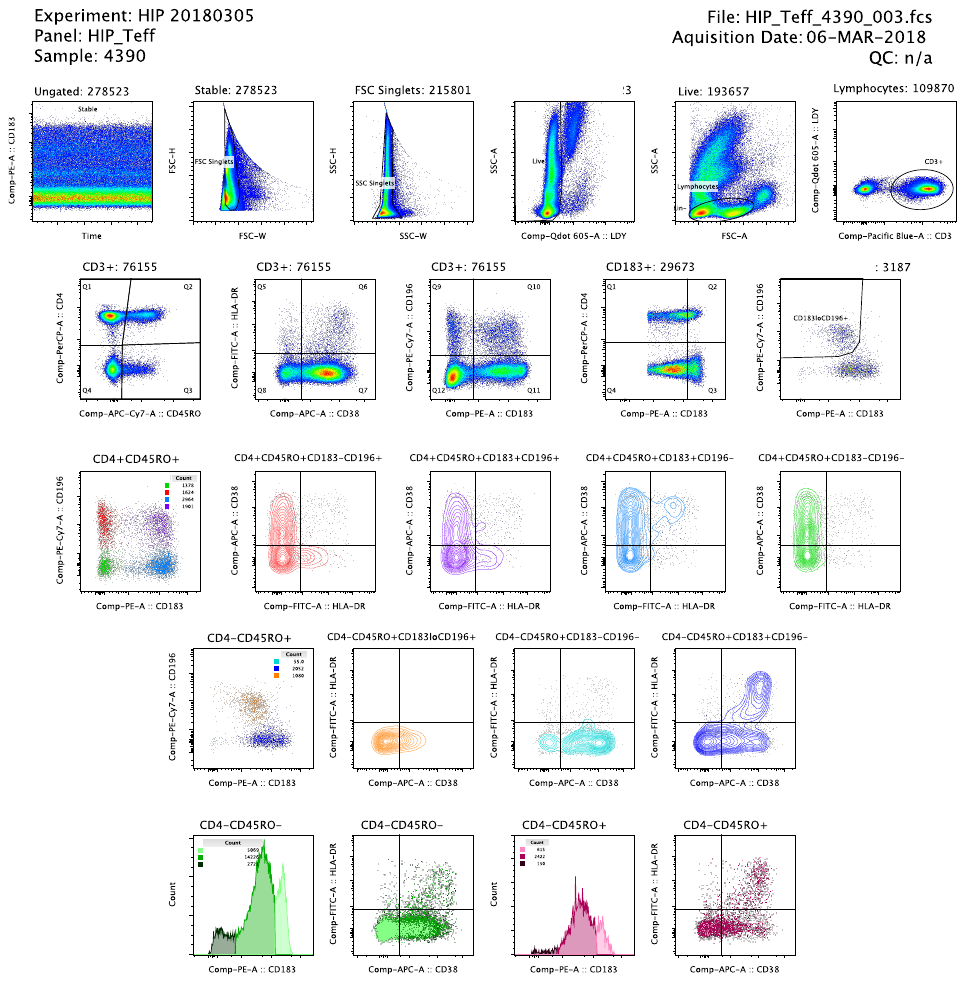
Figure S4. Effector T cell (Teff) panel gating. Gating based on time, forward scatter (FSC), and side scatter (SSC) was performed to select stably acquired singlets. The Qdot605/LDY channel was used to remove autofluorescent granulocytes, then lymphocytes were gated based on FSC and SSC parameters. CD3^+^ T cells were grouped into CD4^+^CD45RO^+^ memory T cells, comprising CD183^-^CD196^+^ Th17 (red), CD183^+^CD196^+^ Th1/17 (purple), CD183^+^CD196^-^ Th1 (blue), and CD183^-^CD196^-^ Th2 (green) subsets. CD3^+^CD4^-^CD45RO^+^ memory CD8 T cells were subset into CXCR3^lo^CCR6^+^ (orange), CXCR3^-^CCR6^-^ (light blue), and CXCR3^+^CCR6^-^ (dark blue) populations. The previous seven cell subsets were further divided into HLA-DR^+^ and HLA-DR^+^CD38^+^ portions. CD3^+^CD4^-^CD45RO^-^ naïve CD8 T cells were gated to CXCR3^-^ (dark green), CXCR3^lo^ (green), CXCR3^hi^ (light green) subsets. CD3^+^CD4^-^CD45RO^+^ memory CD8 T cell populations shown as CXCR3^-^ (dark purple), CXCR3^lo^ (purple), and CXCR3^hi^ (pink).

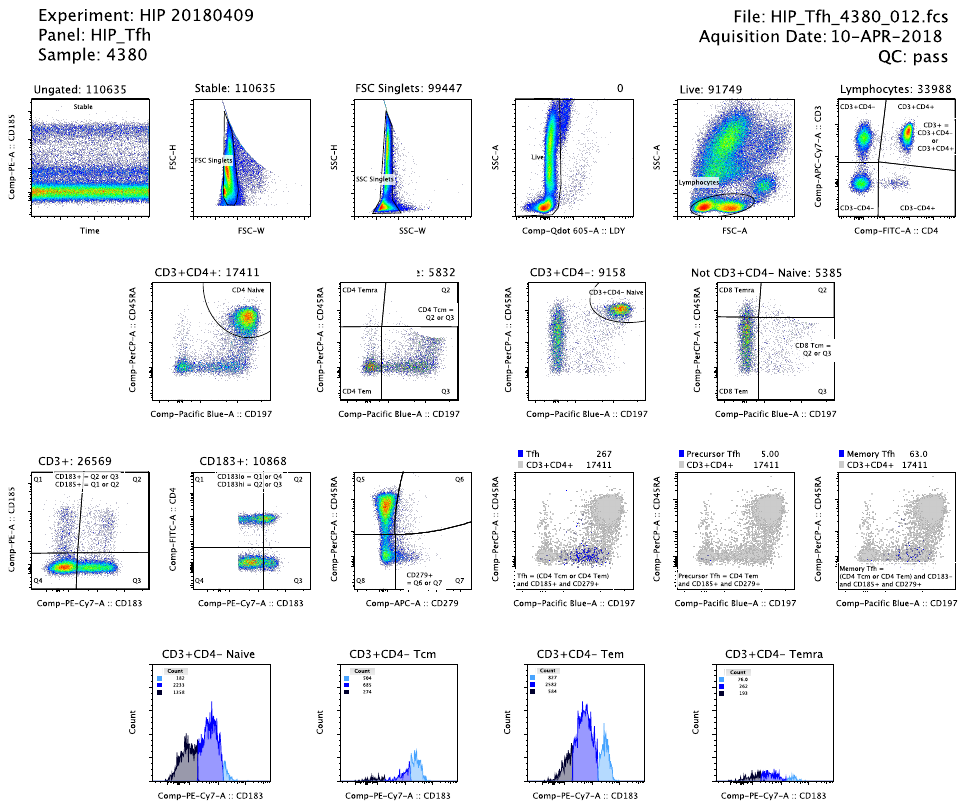
Figure S5. T follicular helper cell (Tfh) panel gating. Gating based on time, forward scatter (FSC), and side scatter (SSC) was performed to select stably acquired singlets. The Qdot605/LDY channel was used to remove autofluorescent granulocytes, then lymphocytes were gated based on FSC and SSC parameters. CD3^+^CD4^+^ and CD3^+^CD4^-^ T cells were grouped into naïve, Tcm, Tem, and Temra based on CD45RA and CD197 expression (see Figure S3). PD-1/CD279^+^, CXCR3/CD183^+^, and CXCR5/CD185^+^ populations were gated from the previously defined CD4 and CD8 T cell subsets. CD4 T cells were gated to CD45RA^-^CXCR5^+^PD-1 ^+^ T follicular helper (Tfh). Tfh were subset into CCR7/CD197^-^ precursor Tfh, CCR7/CD197^+^CD183/CXCR3^-^ memory Tfh, and CXCR3/CD183^+^ Tfh populations. Additionally, CXCR3/CD183^-^ (dark blue), CXCR3/CD183^lo^ (blue), and CXCR3/CD183^hi^ (light blue) populations were determined based on histograms of marker expression as shown, equivalent to method in Fig.S4.

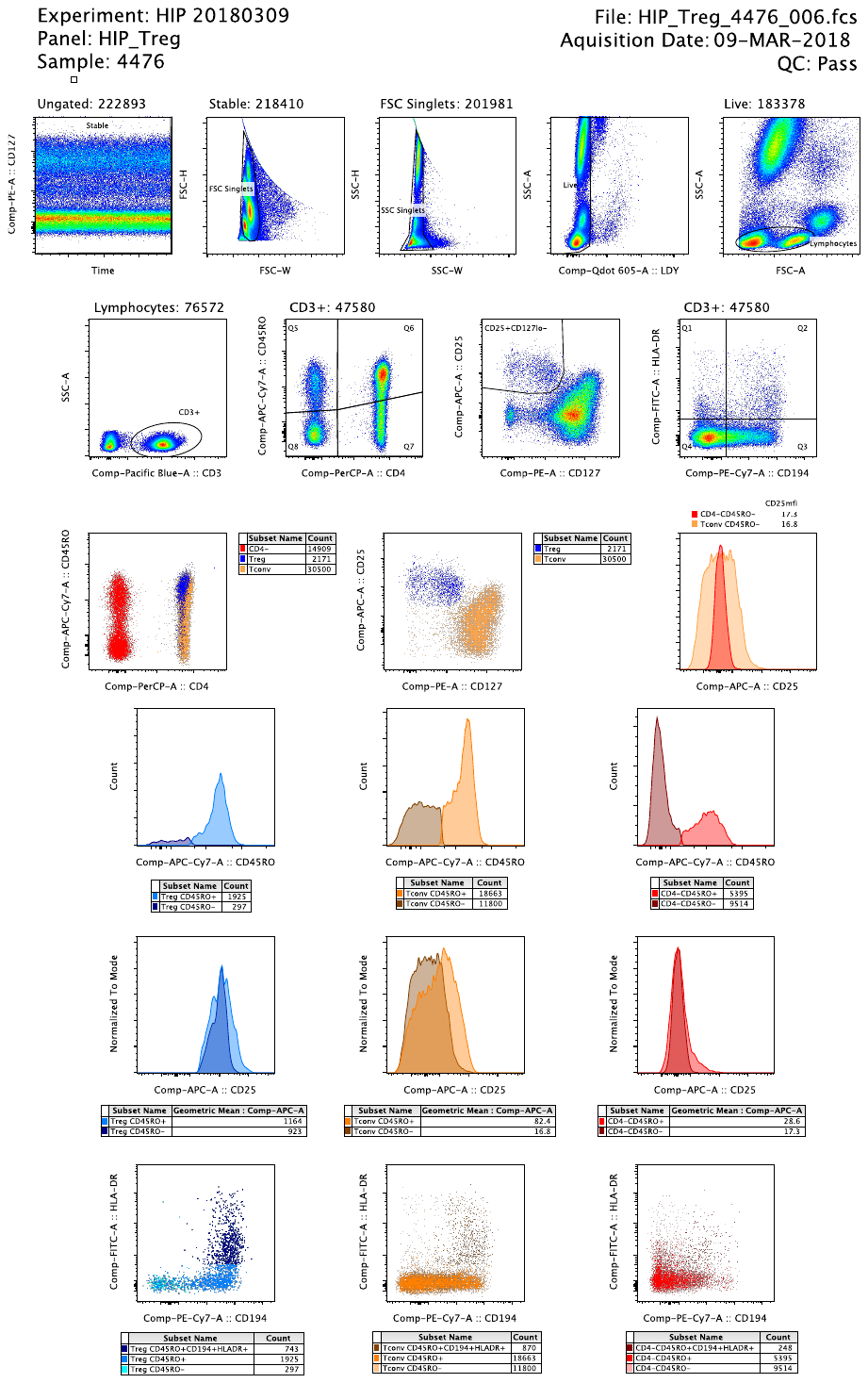

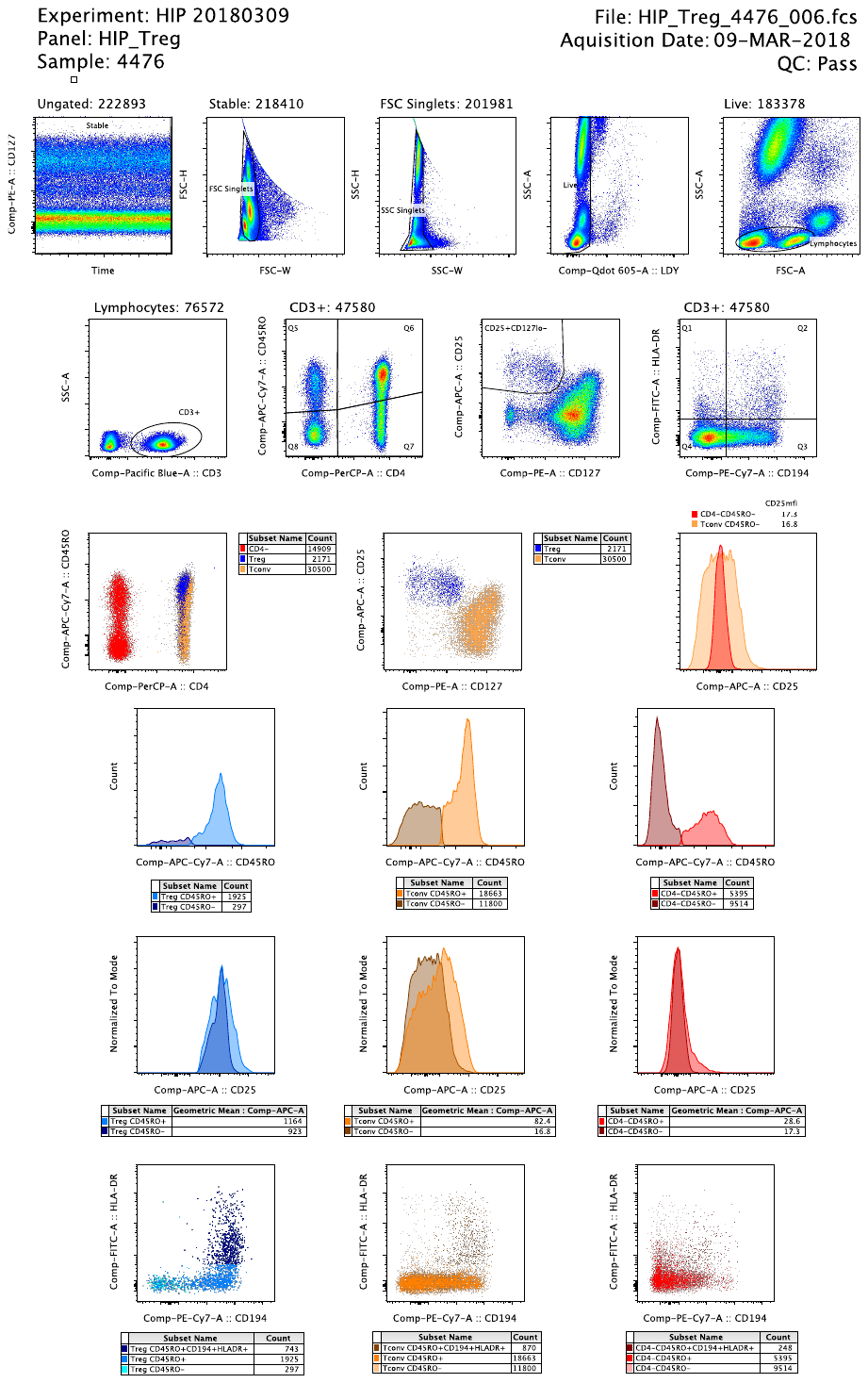

Figure S6. Regulatory T cell (Treg) panel gating. Gating based on time, forward scatter (FSC), and side scatter (SSC) was performed to select stably acquired singlets. The Qdot605/LDY channel was used to remove autofluorescent granulocytes, then lymphocytes were gated based on FSC and SSC parameters. Bottom left: CD3^+^ T cells were grouped into CD4^+^CD25^hi^CD127^lo/-^ Treg (blue), CD4^+^ Tconv as non-Treg (orange), and CD4^-^ to denote CD8 T cells (red). Bottom middle: histogram of CD25 expression on CD3^+^CD4^-^CD45RO^-^ naïve CD8 (red, bottom middle) and CD45RO^-^ naïve Tconv (orange) demonstrates compensation strategy. Right top/middle: CD45RO and CD25 expression on CD45RO^+^ memory Treg (light blue), CD45RO^-^ naïve Treg (blue), CD45RO^+^ memory Tconv (orange), CD45RO^-^ naïve Tconv (brown), CD3^+^CD4^-^CD45RO^+^ memory CD8 (red), and CD3^+^CD4^-^CD45RO^-^ naïve CD8 (dark red) T cells. Right bottom: CCR4/CD194 and HLA-DR expression on CD45RO^+^CCR4^+^HLA-DR^+^ activated memory Treg (dark blue), CD45RO^+^ memory Treg (blue), CD45RO^-^ naïve Treg (light blue), CD45RO^+^CCR4^+^HLA-DR^+^ Tconv/memory Th2 (brown), CD45RO^+^ memory Tconv (orange), CD45RO^-^ naïve Tconv (light orange), CD45RO^+^CCR4^+^HLA-DR^+^ CD8 (dark red), CD4^-^CD45RO^+^ memory CD8 (red), and CD45RO^-^ naïve CD8 (pink) T cells.

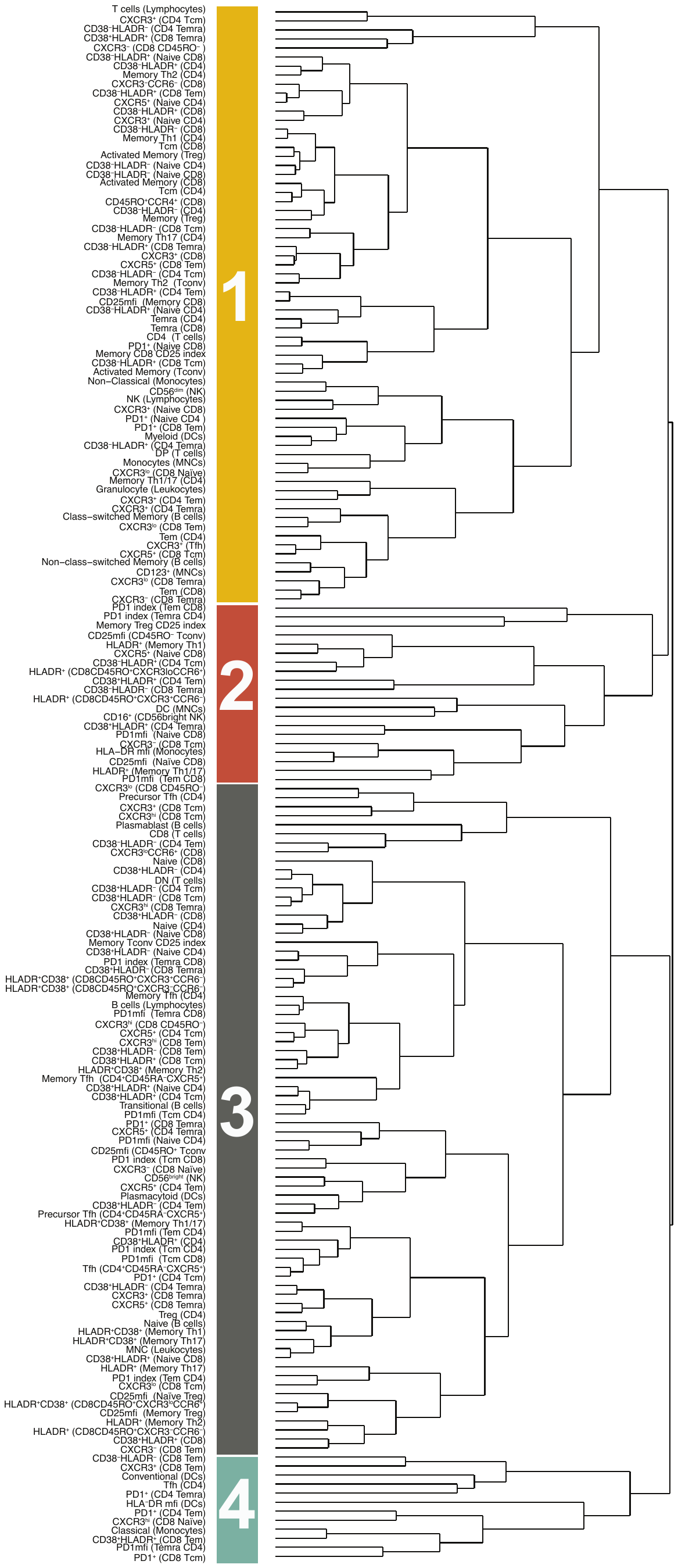

Figure S7. Dendrogram of phenotype age trajectories. Hierarchical clustering of the fitted phenotypic age trajectories resulted in a high-level splitting into four major groups. (See also Figure 2).

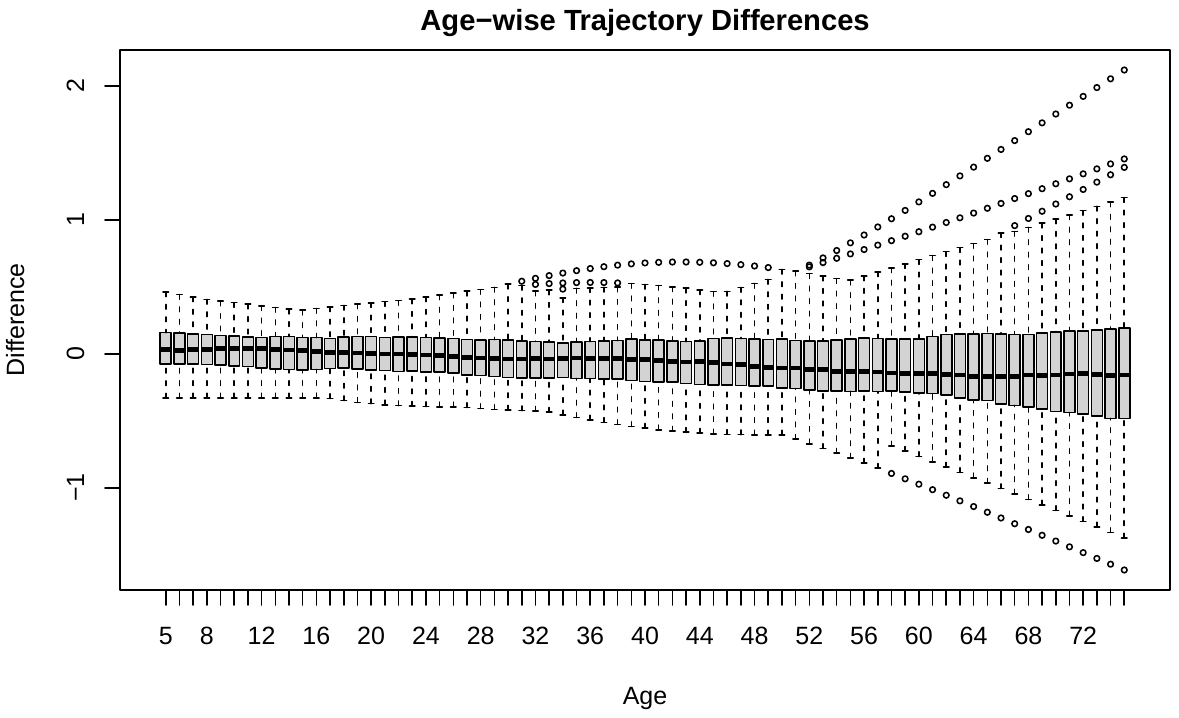

Figure S8. Age-wise trajectory differences. The differences in the estimated trajectory for T1D versus AAb- individuals for each phenotype are shown as boxplots for ages 5-75. (See also Figure 2).

­­
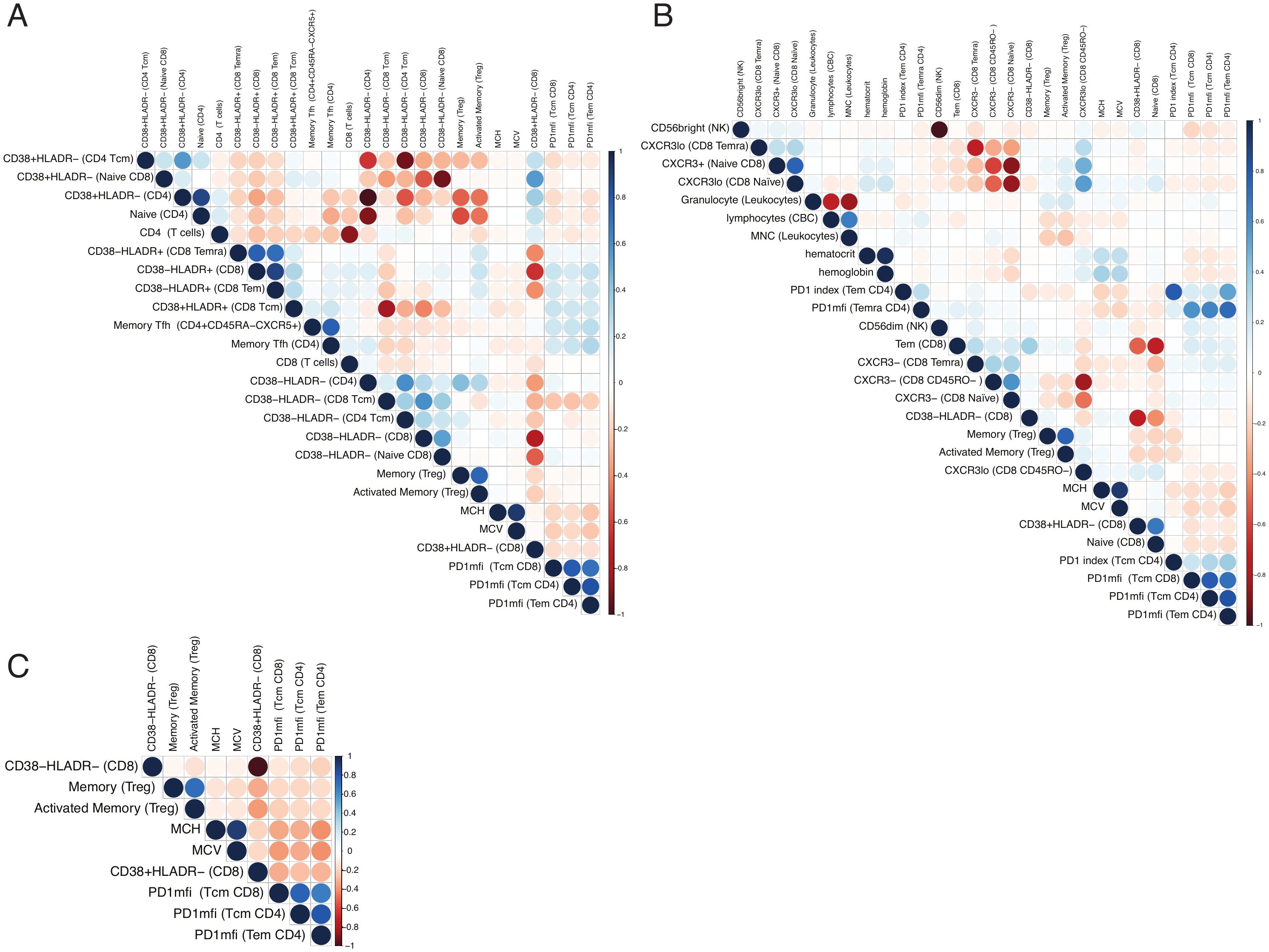

Figure S9. Correlations among significant age-corrected phenotypes. (A) Pairwise Spearman correlation of age-corrected phenotypes that are significantly associated with age and have an absolute correlation larger than 0.7 are shown. Positive correlations are displayed as various shades of blue and negative correlations are indicated by shades of red. (B) Similar to A for phenotypes significantly associated with T1D. (C) Similar to A for phenotypes significant in both age and T1D. (See also Figures 4-5).

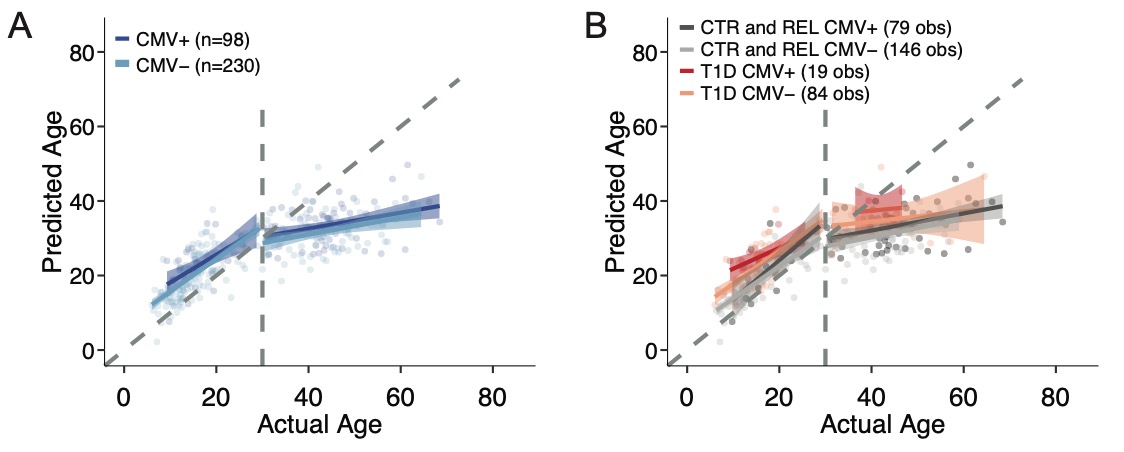

Figure S10. Immune aging model with CMV status. (A) The immunological predicted age is shown for all T1D and AAb- individuals combined. Points are colored based on CMV status with CMV positive in dark blue and CMV negative in light blue. The relationship between predicted age with chronological age is shown using a piece-wise regression model with a break at chronological age 30. (B) Similar in structure to panel A, but additionally split by cohort. Gray represents AAb- individuals (CTR and REL combined) with CMV+ AAb- in dark gray and CMV- AAb- in light gray, while T1D are in red with CMV+ T1D in dark red vs CMV- T1D in pink. (See also Figure 3).

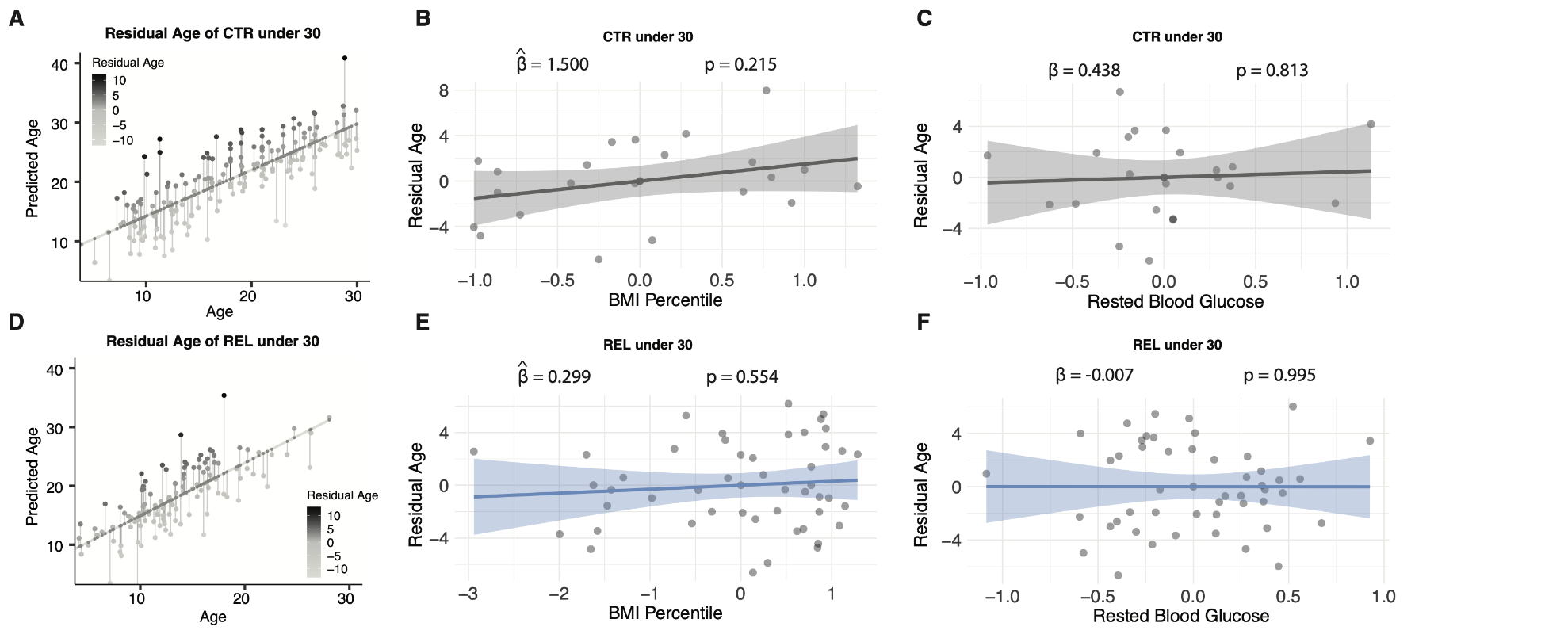

Figure S11. Residual age analysis in AAb- CTR and REL under 30 years of age. (A) Residual immunological age is calculated as the residual from a linear regression of predicted age and chronological age for controls. (B) The partial regression plot of residual age and BMI percentile is shown along with the standardized regression coefficient and p-value. (C) Similar in structure to B for the association between residual age and rested blood glucose. (D) Similar in structure to panel A, but for AAb- REL. (E-F) similar in structure to panels B and C for AAb- REL. (See also Figure 3).

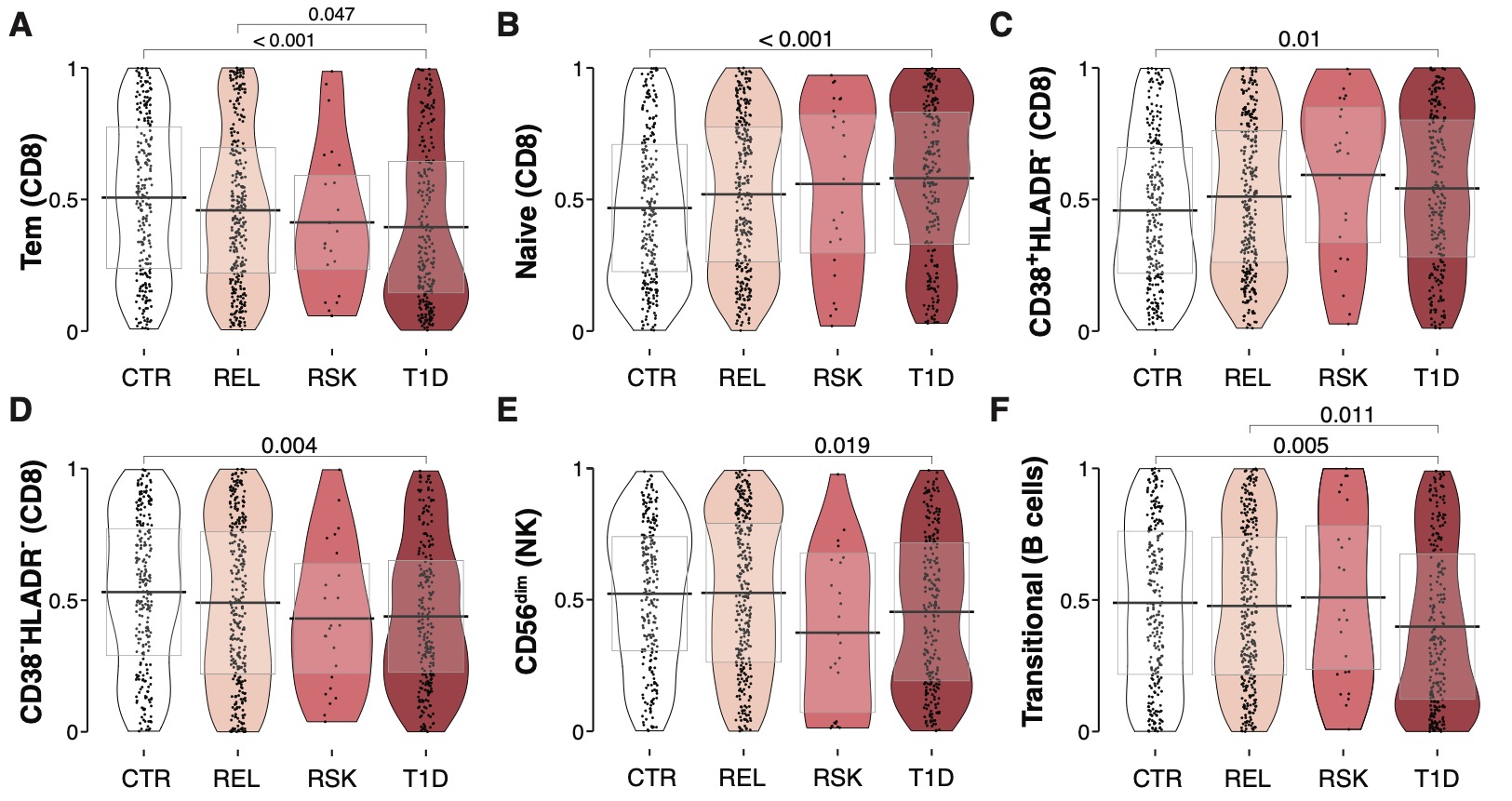

Figure S12. Adaptive immune compartment findings confirming prior literature. Immune subsets from AAb- CTR, AAb- REL, ≥2 AAb+ RSK, and T1D individuals. Significant p-values shown on graph. Testing was done using a Kruskal Wallis test with a post-hoc Dunn’s test and Benjamini-Hochberg multiplicity adjustment. The y-axis represents age-adjusted quantile values.

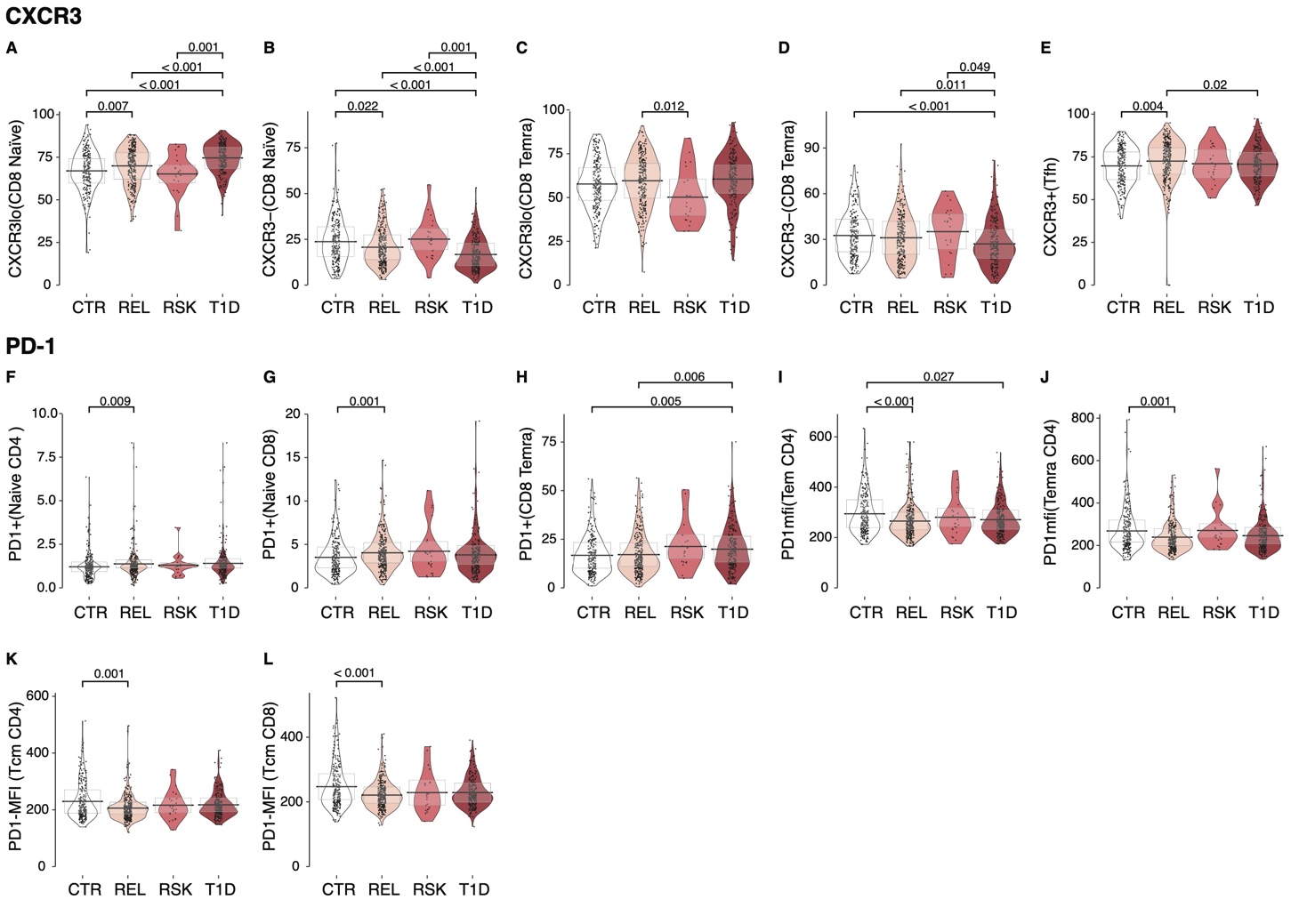
Figure S13. Raw data shows CXCR3 and PD-1 expression is increased on naïve T cells but decreased on memory T cell subsets in AAb- CTR, REL, ≥2 AAb+ and T1D individuals. (A-D) CXCR3 low or negative proportions on CD8 subsets, (E) CXCR3 positive proportions of CD4 Tfh cells, (F-H) PD-1^+^ and (I-L) PD-1 MFI for each phenotype shown from AAb- CTR, AAb- REL, ≥2 AAb+ RSK, and T1D individuals. Significant p-values shown on graph. Testing was done using a Kruskal Wallis test with a post-hoc Dunn’s test and Benjamini-Hochberg multiplicity adjustment (See also Figure 6).

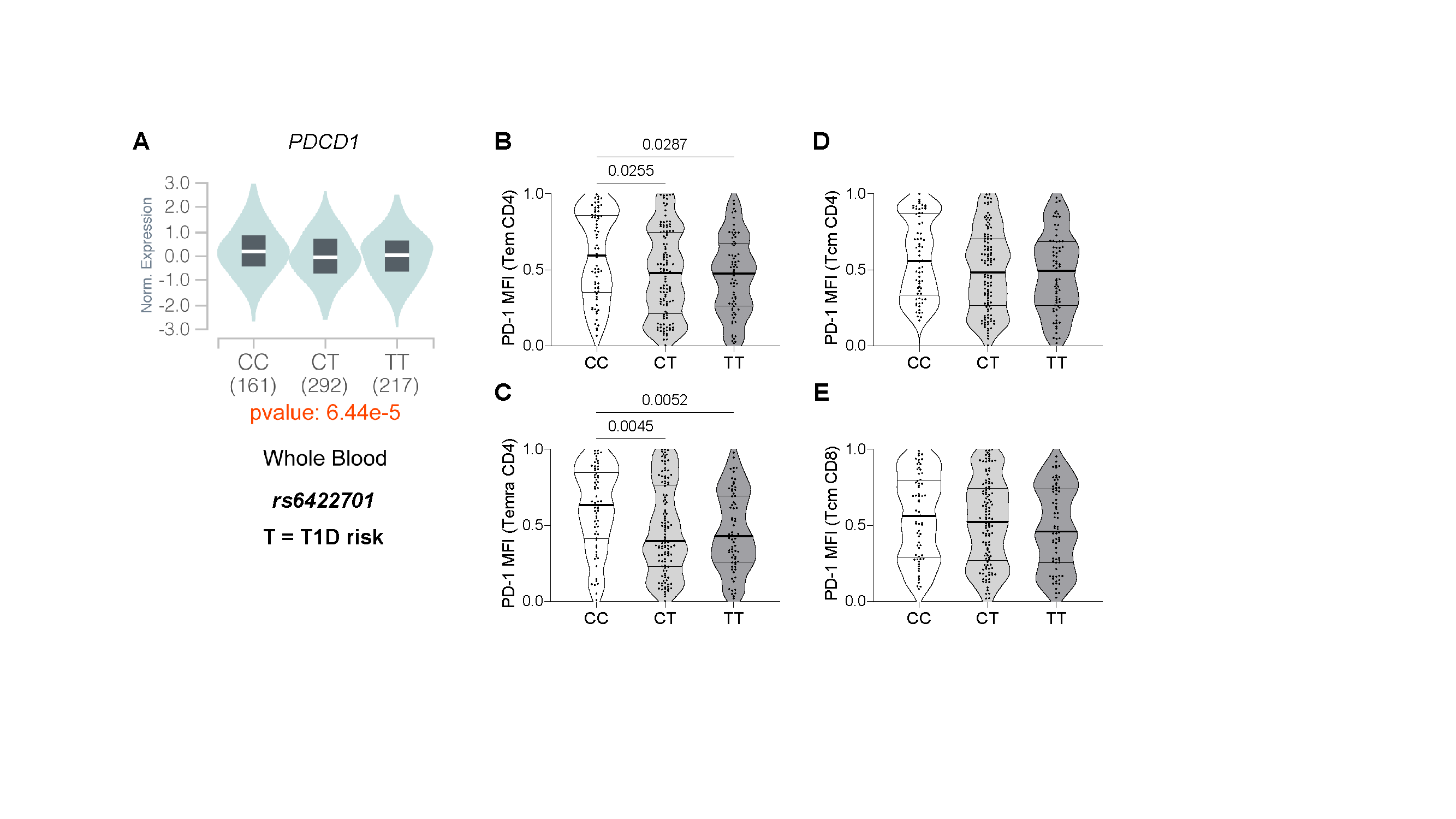

Figure S14. *PDCD1* expression quantitative trait locus (eQTL), rs6422701, associates with PD-1 mRNA and protein expression. (A) Genotype-Tissue Expression project (GTEx) data shows normalized *PDCD1* mRNA expression in whole blood differs according to genotype at rs6422701. In our cohort, the T allele of rs6422701 was overrepresented in the T1D group (See Table S4). Age-adjusted quantile values of PD-1 MFI on (B) CD4 Tem, (C) CD4 Temra, (D) CD4 Tcm, and (E) CD8 Tcm of AAb- CTR and AAb- REL according to rs6422701 genotype. Significant p-values shown on graph. Testing was done using a Kruskal Wallis test with a post-hoc Dunn’s test.
